# Single-Cell multiomics reveals ENL mutation perturbs kidney developmental trajectory by rewiring gene regulatory landscape

**DOI:** 10.1101/2024.05.09.591709

**Authors:** Lele Song, Qinglan Li, Lingbo Xia, Arushi Sahay, Qi Qiu, Yuanyuan Li, Haitao Li, Kotaro Sasaki, Katalin Susztak, Hao Wu, Liling Wan

**Author notes:** Correspondence should be addressed to L.W. These authors contributed equally (Lele Song, Qinglan Li).

## Abstract

Cell differentiation during organogenesis relies on precise epigenetic and transcriptional control. Disruptions to this regulation can result in developmental abnormalities and malignancies, yet the underlying mechanisms are not well understood. Wilms tumors, a type of embryonal tumor closely linked to disrupted organogenesis, harbor mutations in epigenetic regulators in 30-50% of cases. However, the role of these regulators in kidney development and pathogenesis remains unexplored. By integrating mouse modeling, histological characterizations, and single-cell transcriptomics and chromatin accessibility profiling, we show that a Wilms tumor-associated mutation in the chromatin reader protein ENL disrupts kidney development trajectory by rewiring the gene regulatory landscape. Specifically, the mutant ENL promotes the commitment of nephron progenitors while simultaneously restricting their differentiation by dysregulating key transcription factor regulons, particularly the *HOX* clusters. It also induces the emergence of abnormal progenitor cells that lose their chromatin identity associated with kidney specification. Furthermore, the mutant ENL might modulate stroma-nephron interactions via paracrine Wnt signaling. These multifaceted effects caused by the mutation result in severe developmental defects in the kidney and early postnatal mortality in mice. Notably, transient inhibition of the histone acetylation binding activity of mutant ENL with a small molecule displaces transcriptional condensates formed by mutant ENL from target genes, abolishes its gene activation function, and restores developmental defects in mice. This work provides new insights into how mutations in epigenetic regulators can alter the gene regulatory landscape to disrupt kidney developmental programs at single-cell resolution *in vivo*. It also offers a proof-of-concept for the use of epigenetics-targeted agents to rectify developmental defects.

## Introduction

Normal cellular differentiation is a tightly regulated process that relies on finely tuned epigenetic and transcriptional control^1–3^. Germline or somatic mutations affecting epigenetic/transcriptional regulators can disrupt these mechanisms, leading to developmental disorders and malignancies^2,4,5^. This phenomenon is particularly evident in pediatric cancers^6–8^, where the roots are suspected to be intertwined with aberrant developmental trajectories. However, in most tissue types, the dynamic chromatin and gene regulatory events that orchestrate cell fate determination and how alterations in these events impact normal development and drive the diseased state remain unclear. Unraveling such insights has the potential to identify vulnerable cell types or developmental states as the disease origins and pave the way for precision medicine.

Wilms tumor, the most common pediatric kidney tumor^9,10^, is linked to disrupted embryonic kidney development^10–12^. As such, it serves as a paradigm for understanding the interplay between development and tumorigenesis. The mammalian kidney emerges from the intermediate mesoderm through reciprocal interactions between two tissues, the ureteric bud (UB) and the metanephric mesenchyme^13–15^. At the initiation of nephrogenesis, signals from the mesenchyme induce reiterative branching of the UB. UB-derived signals in turn induce a subset of nephron progenitor cells (NPCs) located in the cap mesenchyme (CM) surrounding the UB tip to commit and undergo mesenchymal-to-epithelial transition (MET). This transition gives rise to an intermediate condensed structure known as the peritubular aggregate (PA), which then progresses into an epithelial structure termed the renal vesicle (RV). Subsequent segmentation and elongation of the renal vesicle give rise to a variety of epithelial nephron structures, including glomerular podocytes, proximal tubules (PT), loops of Henle (LOH), and distal tubules (DT), while the UB becomes the collecting duct^13–15^ (**Figure 1a**). The metanephric mesenchyme also gives rise to the renal stroma^16^, which plays an important role in proper differentiation of the UB and nephron^17–19^.

**Figure 1.**
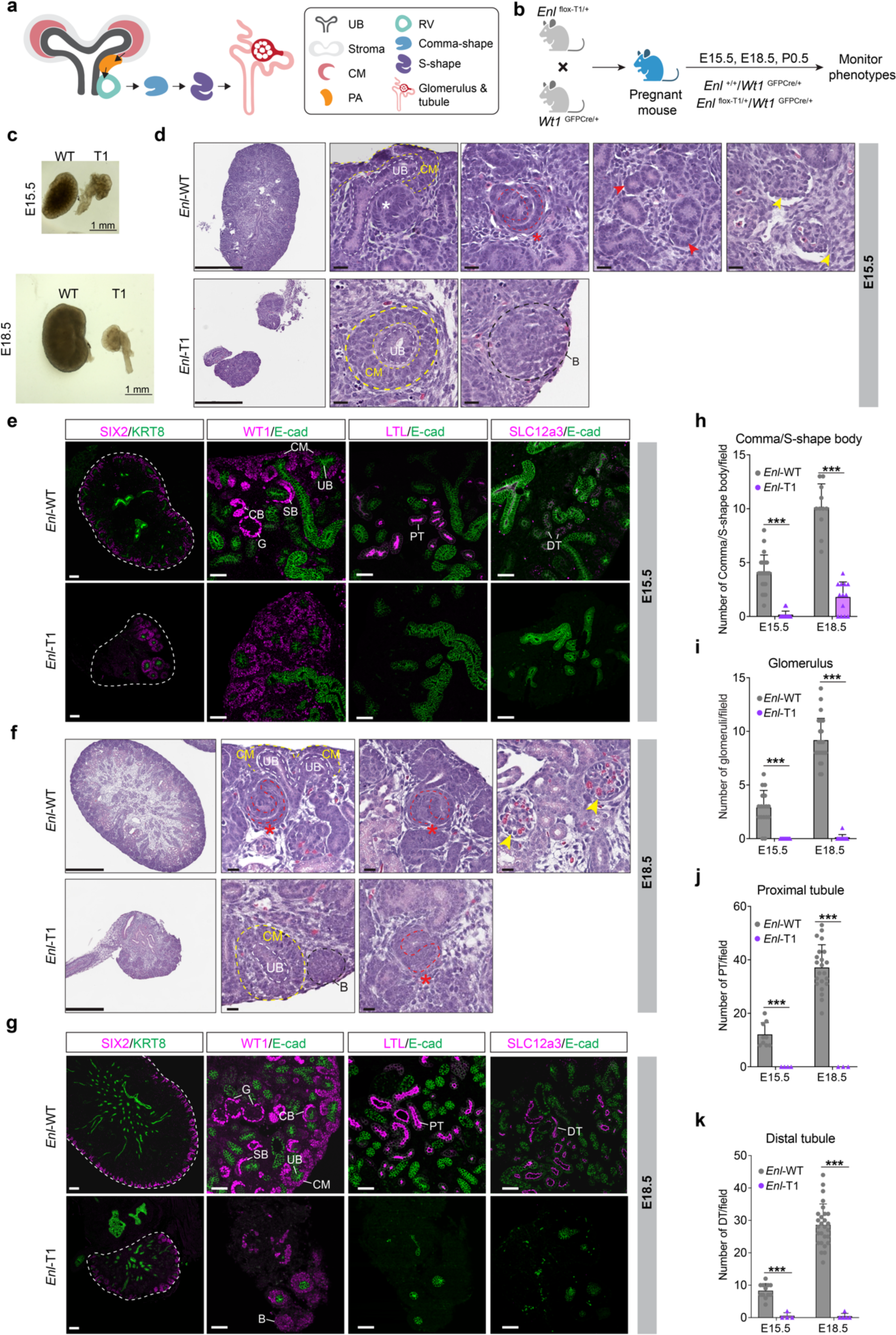
Heterozygous expression of mutant ENL disrupts embryonic kidney development and leads to postnatal mortality in mice. **a**, Schematic of nephrogenesis. Cap mesenchyme (CM), ureteric bud (UB), peritubular aggregate (PA), renal vesical (RV). **b**, Schematic of the mouse breeding and experimental strategy. **c**, Brightfield images of the E15.5 (top) and E18.5 (bottom) kidneys. Scale bar, 1 mm. **d**, **f**, Hematoxylin and eosin-stained sections showing the histology of E15.5 (**d**) and E18.5 (**f**) kidneys from *Enl*-WT and *Enl*-T1 embryos. The red star indicates S-shape body, the red arrows indicate tubules, the black dashed circle indicates a region of blastema-like structure (B), and the yellow arrows indicate glomeruli structures. Scale bar in the first column images, 250 µm (**d**) and 500 µm (**f**); scale bar in the zoom-in images, 20 µm. **e**, **g**, Immunostaining for SIX2, KRT8, WT1, E-cadherin (E-cad), LTL, and SLC12a3 at E15.5 (**e**) and E18.5 (**g**) kidney sections. Scale bar in the first column images, 150 µm (**e**) and 100 µm (**g**); scale bar in the zoom-in images, 50 µm. SB, S-shape body; CB, Comma-shape body; G, glomerulus; PT, proximal tubule; DT, distal tubule. **h-k**, Quantification for the numbers of nephron structures per field. **h**, Comma/S-shape body (E15.5, *n* = 9 WT and 10 T1 kidneys; E18.5, *n* = 6 WT and 6 T1 kidneys); **i**, glomerulus (E15.5, *n* = 7 WT and 9 T1 kidneys; E18.5, *n* = 9 WT and 7 T1 kidneys); **j**, proximal tubule (E15.5, *n* = 7 WT and 4 T1 kidneys; E18.5, *n* = 7 WT and 3 T1 kidneys); **k**, distal tubule (E15.5, *n* = 7 WT and 4 T1 kidneys; E18.5, *n* = 6 WT and 5 T1 kidneys). One dot indicates the number of indicated structures per field. Date represent mean ± s.d.; two-tailed unpaired Student’s *t*-test, \*\*\**P* < 0.001.

Histologically, Wilms tumor closely resembles the embryonic kidney and is often marked by rudimentary structures^12^. Known molecular drivers of Wilms tumor commonly involve disruptions to key transcription factors (TFs) (e.g., *WT1*, *SIX2*) and signaling proteins (e.g., *IGF2*, *WTX*, β-catenin) crucial for nephrogenesis^5,20^. To date, Wilms tumor studies have primarily focused on phenotypic characterization of mouse models for a few established molecular players^21–23^, leaving the underlying cellular and molecular mechanism incompletely understood. In addition, the precise causes of two-thirds of Wilms tumor remain unclear. Recent genomic characterization of high-risk Wilms tumor has revealed previously unidentified mutations in epigenetic regulators in 30-50% of cases^20^, underscoring unexplored roles of epigenetics dysregulation in this disease. Furthermore, while recent single-cell profiling studies on both normal kidneys and Wilms tumors^24–28^ have offered valuable insights into kidney development and support a fetal origin for Wilms tumor, significant gaps remain in our understanding of how perturbations in chromatin mechanisms by disease mutations impact cell fate determination during kidney development and pathogenesis, particularly at single-cell resolution.

Our current study aims to address these fundamental questions by focusing on the epigenetic reader protein Eleven-nineteen leukemia (ENL). ENL, also known as MLLT1, exerts its function by binding to histone acylation through its conserved YEATS (Yaf9, ENL, AF9, Taf14, Sas5) domain and recruiting elongation factors to promote transcription^29,30^. ENL plays a crucial role in maintaining subsets of acute myeloid leukemia (AML)^29,30^. Our group has recently developed a small-molecule inhibitor designed to target ENL’s acyl-binding activity, which has shown promising efficacy against AML in animal models^31^. More recently, a series of hotspot mutations within ENL’s YEATS domain have been identified in 5-9% of Wilms tumor patients^32^, making ENL the most frequently mutated epigenetic regulator in this disease^20,32^. Wilms tumors harboring ENL mutations often display intralobular nephrogenic rests, which stem from early kidney development and are associated with a high risk^20,32^. Previous studies in the human embryonic kidney cell line HEK293 have revealed that these mutations confer gain-of-function properties to ENL, enabling it to drive aberrant transcription activation through the formation of condensates at specific target gene loci^33,34^. Moreover, introduction of these ENL mutations into mouse embryonic stem cells has led to the formation of Wilms tumor-like blastema structures in an *in vitro*-directed differentiation assay^35^, suggesting their potential biological significance. However, the precise functions of ENL mutations on kidney development and tumorigenesis *in vivo*, as well as the underlying mechanisms, have remained unknown.

Here, by integrating genetic mouse modeling, histological characterizations, and single-cell transcriptomics and chromatin accessibility profiling, we reveal ENL mutation-induced alterations in cellular composition, differentiation trajectories, and gene regulatory landscapes during the development of the mouse kidney. These cellular and molecular alterations result in impaired nephrogenesis and postnatal mortality in mice. Furthermore, we demonstrate that transient inhibition of the acyl-binding activity of mutant ENL can effectively abolish its chromatin function in cellular systems and more importantly, rescue transcriptomic and developmental defects induced by the mutant in embryonic kidneys *in vivo*. This study provides functional and mechanistic insights into the impact of Wilms tumor-associated ENL mutations on kidney development and offers a proof-of-concept for the use of epigenetics-targeted agents in the correction of developmental defects.

## Results

### Heterozygous expression of mutant ENL disrupts embryonic kidney development and leads to early postnatal mortality in mice

To investigate the role of ENL mutants in nephrogenesis, we generated a conditional knock-in mouse model for the most prevalent *ENL* mutation found in Wilms tumor^32^ (p.117_118insNHL, referred to as ENL-T1) using an inversion strategy (**Figure S1a**). Before induction of Cre recombinase activity, the targeted allele is expressed as *Enl*-WT (**Figure S1a**). After two steps of Cre-mediated recombination, the inverted exon 4 containing the T1 mutation is flipped to the correct direction and expressed, and WT exon 4 is excised, thus leading to the expression of the targeted allele as *Enl*-T1 (**Figure S1a**). Genotyping PCR was used to successfully distinguish the wildtype and the targeted allele before Cre-mediated recombination (**Figure S1b**). To induce *Enl*-T1 expression in the developing kidney, we crossed *Enl* ^flox-T1/+^ mice with *Wt1*^GFPCre/+^ mice (**Figure 1b**) to generate *Enl* ^flox-T1/+^*Wt1*^GFPCre/+^ (hereafter referred to *Enl*-T1) and *Enl*^+/+^*Wt1*^GFPCre/+^ (hereafter referred to *Enl*-WT) offspring. *Wt1* (Wilms tumor 1) is a transcription factor expressed within the intermediate mesoderm^36,37^, the origin of the metanephric kidney^12^. The *Wt1*^GFPCre^ knock-in allele in mice expresses an EGFPCre fusion protein from the *Wt1* promoter/enhancer elements and concomitantly inactivates the endogenous *Wt1* gene^38^. Given the well-characterized expression pattern and function of *Wt1* in kidney development, the *Wt1*^GFPCre^ strain is widely used for genetic studies in kidney biology and diseases^21^. To validate *Enl*-T1 expression following Cre-mediated recombination, mRNA was extracted from E15.5 *Enl*-WT and *Enl*-T1 kidneys for reverse transcription, PCR amplification, and next-generation sequencing to determine the relative abundance of *Enl*-WT and *Enl*-T1 mRNA levels (**Figure S1c**). As expected, only *Enl*-WT mRNA was present in *Enl*-WT kidneys. In *Enl*-T1 kidneys, 58% and 41% of the sequencing reads corresponded to *Enl*-WT and *Enl*-T1 cDNA, respectively (**Figure S1d**). These results confirm the successful induction of *Enl*-T1 and demonstrate comparable expression levels of the *Enl*-WT and *Enl*-T1 alleles in *Enl*^flox-T1/+^/Wt1^GFPCre/+^ kidneys.

While *Enl*-T1 and *Enl*-WT pups derived from the breeding scheme (**Figure 1b**) were born at a mendelian ratio, all *Enl*-T1 newborns exhibited early postnatal mortality (**Figure S1e**). Further examination of embryonic kidneys harvested at embryonic day 15.5 and 18.5 (E15.5 and E18.5) revealed markedly reduced size of *Enl*-T1 kidneys compared to their WT counterparts (**Figure 1c**). Histologically, *Enl*-WT kidney harbored expected structures characteristic of the developing kidney including ureteric buds (UB) invading the cap mesenchyme (CM), intermediate differentiating nephron structures (comma/S-shape bodies), and fully differentiated nephron structures (tubules and glomeruli) (**Figure 1d**, **f**). These results, aligning with earlier findings^39^, confirm that the loss of one allele of *Wt1* has minimal impacts on embryonic kidney development, supporting these mice as suitable controls for our study. In *Enl*-T1 kidneys, although the CM and UB structures were discernible, the CM was characterized as multilayers of cells enveloping a non-branching UB (**Figure 1d**, **f**). In addition, some *Enl*-T1 kidneys exhibited proliferative structures with elevated Vimentin expression that morphologically resembled undifferentiated blastema components observed in Wilms tumors^12^ (**Figure 1d**, **f, Figure S1f**). The CM abnormality in *Enl*-T1 kidneys was further validated through immunostaining for SIX2 (a marker for NPCs)^40^, KRT8 (a marker for the ureter epithelium)^41^, WT1 (expressed in NPCs and podocytes)^36^, and E-cadherin (epithelial cells)^42^ (**Figure 1e, g**). Moreover, *Enl*-T1 kidneys displayed fewer differentiating/differentiated structures such as the comma/S-shape bodies (**Figure 1d-h**), glomeruli (**Figure 1d-g, i**), and elongating tubules including the PT (LTL^+^) and DT (SLC12a3^+^) (**Figure 1d-g, j, k**). These defects in the developing kidney of *Enl*-T1 mice persisted postnatally (**Figure S1g-l**), which likely explains the observed postnatal mortality (**Figure S1e**). Thus, heterozygous expression of *Enl*-T1 initiated in *Wt1*^+^ metanephric mesenchyme precursors substantially disrupts kidney development primarily through a gain of aberrant CM/UB structures and a reduction in differentiated nephron structures. This perturbation ultimately results in kidney dysfunction and postnatal lethality in mice.

Given that *Wt1*^GFPCre^ induces *Enl*-T1 expression in both nephron and stroma progenitors^43,44^, we next asked whether *Enl*-T1 expression in both lineages contributes to the developmental defects observed in *Enl* ^flox-T1/+^*Wt1*^GFPCre/+^ mice. To this end, we crossed *Enl*^flox-T1/+^ mice with *Six2*^GFPCre/+^ mice to induce *Enl*-T1 expression specifically in *Six2*^+^ NPCs and their progeny^40^ (**Figure S2a)**. Similar to findings with the *Wt1*^GFPCre/+^ strain, *Enl*-T1 expression caused early postnatal lethality in the *Six2*^GFPCre/+^ strain (**Figure S2b**). However, the phenotypic manifestations of *Enl*-T1 expression differed between the two Cre strains. Specifically, *Enl*-T1 expression in *Six2*^+^ NPCs had minimal impact on the overall kidney size and tubular structures, but significantly affected the development and maturation of glomeruli in embryonic (E15.5 and E18.5) (**Figure S2c-f**) and neonatal (P0.5) (**Figure S2g, h**) kidneys, as indicated by reduced numbers as well as shrunken and fragmented glomeruli. These results suggest that expression of *Enl*-T1 can impair kidney development through its effects in both nephron and stroma compartments.

### Single-cell transcriptomics reveals altered cellular composition and differentiation trajectories in *Enl*-mutant kidneys

To dissect the mechanism by which mutant ENL perturbs embryonic kidney development, we isolated and sequenced a total of 10, 000 cells from whole kidney suspensions derived from 2 *Enl*-WT and 4 *Enl*-T1 E15.5 embryos. Following stringent filtration, 9376 individual cells (5232 for WT and 4144 for T1) were retained for further analysis (**Figure S3a-c**). We first focused on developing a single-cell transcriptomics map of *Enl*-WT kidneys. Unbiased clustering coupled with the expression patterns of lineage-specific marker genes identified four major lineage compartments: nephron (*Six2^+^* and *Cited1^+^*), stroma (*Pdgfra^+^* and *Col1a1^+^*), UB/ureteric epithelium (UE) (*Ret^+^* and *Calb1^+^*), and endothelial cells (*Pecam1^+^* and *Cdh5^+^*) (**Figure S3d**, **e**)^45,46^. Within the nephron, cells were further grouped into 10 clusters (**Figure S3f**) representing various cell types, including nephron progenitors (C0/1/4: *Six2*^+^), distal tubule precursor (C6-DT pre: *Gata3*^+^, *Sim1*^+^), distal tubule (C8-DT: *Gata3*^+^), LOH precursor (C5-LOH pre: *Sim1*^+^), LOH (C9-LOH: *Sl*c12a1^+^), proximal tubule (C7-PT: *Slc34a1*^+^), and podocyte (C3-podo: *Nphs1*^+^, *Mafb*^+^) (**Figure S3g**). Within nephron progenitors, we identified two major cell states: Cluster 0 represents self-renewing NPCs with high expression levels of *Six2* and *Cited1* (C0-NP1: *Six2*^+^/*Cited1*^+^), while clusters 1/4 represent NPCs primed for differentiation (C1/4-NP2: *Six2*^low^*/Wnt4*^+^) that are often found within the pretubular aggregate (**Figure S3g**). In addition, we identified a cluster representing an intermediate cell state (C2-IM: *Cdh6*^+^, *Six2*^-^), which loosely corresponds to cells in the renal vesicle (RV), the precursor of nephron^15^. Developmental trajectory analysis of *Enl*-WT nephrons revealed three distinct differentiation paths originating from NPCs, leading to the formation of podocytes, PT, and DT/LOH, respectively (**Figure S3h**). These results align well with prior characterizations and single cell analyses on normal mouse kidneys^26,28,45,47^. Thus, despite the heterozygosity of *Wt1* in *Enl*-WT kidneys, their cellular composition and differentiation pattern closely resemble those of a typical, healthy kidney.

To assess the consequences of the *ENL* mutation, we analyzed the integrated scRNA-seq datasets from *Enl*-WT and *Enl*-T1 kidneys. Using single-cell transcriptomes of *Enl*-WT cells as a reference map for cell type annotations, we showed that *Enl*-T1 cells were similarly grouped into 4 lineage compartments (**Figure 2a**). A quantitative analysis of the cellular distribution among these compartments revealed a relative decrease in the nephron and an increase in the stroma in *Enl*-T1 kidneys (**Figure 2b**). These observations, coupled with the fact that *Wt1*-cre expression is restricted to the nephron and stroma (**Figure S3i, j**), prompted us to focus our subsequent analyses on the role of mutant ENL in these two lineages.

**Figure 2.**
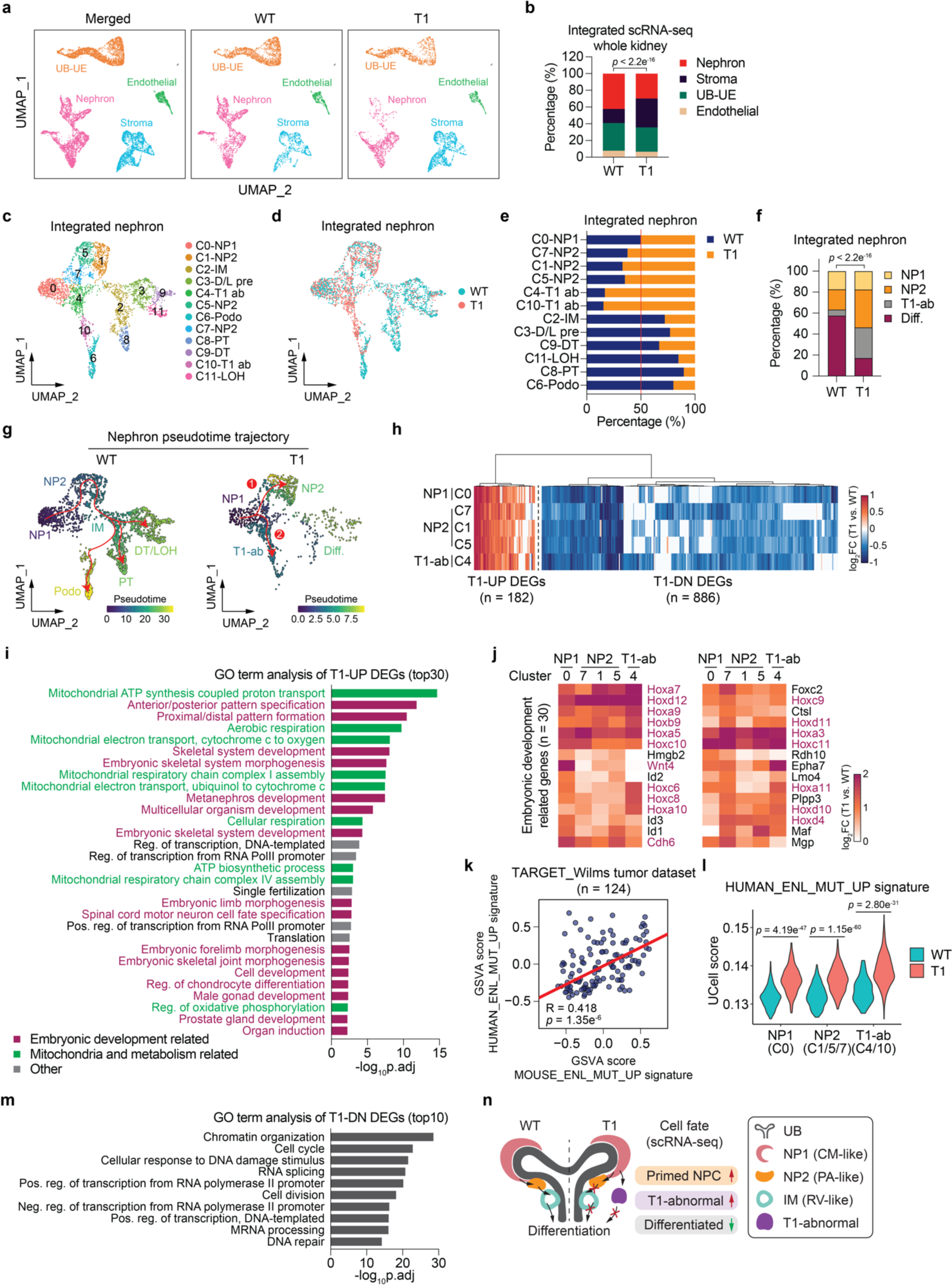
Mutant ENL alters the cellular composition, differentiation trajectories, and gene expression programs in the developing kidney. **a**, UMAP embedding of scRNA-seq data labeled with four main embryonic kidney lineages. UB-UE, ureteric bud-ureteric epithelium. **b**, The percentage of four main embryonic kidney compartments within samples. Chi-Square test *p*-value is shown. **c**, **d**, UMAP embedding of integrated scRNA-seq cells from *Enl*-WT and T1 nephrons. Cells are colored and labeled by annotated cell types (**c**) or samples (**d**). D/L pre, distal tubule/loop-of-Henle precursor; T1 ab, T1-abnormal; Podo, podocyte; LOH, loop-of-Henle. **e**, The relative percentage of cell types between scRNA-seq *Enl*-WT and T1 nephrons. **f**, The percentage of cell types within scRNA-seq *Enl*-WT or T1 nephron. Diff., differentiation, including RV, Podo, PT, D/L pre, DT, and LOH cell types. Chi-Square test *p*-value is shown. **g**, UMAP embedding of *Enl*-WT and T1 scRNA-seq nephron differentiation trajectory. Cells are colored by pseudotime. Trajectories are depicted in red. Two distinct differentiation paths in *Enl*-T1 nephron are highlighted as “1” for the normal path and “2” for the abnormal path. **h**, The fold change of differentially expressed genes (DEGs) between *Enl*-WT and T1 within the indicated nephron cell types. Fold change is scaled by log2. T1-UP, upregulated in *Enl*-T1 cells; T1-DN, downregulated in *Enl*-T1 cells. See Supplementary Tables 2. **i**, **m**, Gene ontology (GO) term analysis for the union T1-UP (**i**) and DN (**m**) DEGs shown in (**h**). Embryonic development-related terms are colored in dark red; mitochondrial and metabolism-related terms are colored in green; others are colored in grey. **j**, The log2 fold change of embryonic development-related T1-UP DEGs identified in (**i**) within the indicated nephron cell types. *Hox, Wnt4, and Cdh6* genes are highlighted in dark red. **k**, Pearson correlation analysis of the Gene Set Variation Analysis (GSVA) scores evaluated by human and mouse ENL_MUT_UP signatures for the Wilms tumor patients from the TARGET dataset. See Supplementary Tables 4. Each dot represents the GSVA scores of one TARGET_Wilms tumor patient. R, Pearson correlation coefficient. **l**, The UCell score evaluated by human ENL_MUT_UP signature for the indicated nephron cell types within *Enl*-WT and T1 datasets, respectively. Wilcoxon rank-sum test *p*-values are shown. **n**, Schematic illustrating the impaired nephrogenesis in the *Enl*-T1 nephron revealed by scRNA-seq.

In the integrated nephron, we identified 12 distinct cell clusters that correspond to 9 different cell types (**Figure 2c**). There was a noticeable depletion of differentiated cell types, such as PT (C8), LOH (C11), DT (C9), and podocytes (C6) in *Enl*-T1 nephrons compared with wildtype. Conversely, undifferentiated cell types, such as NP2 (C1/5/7), displayed a slight enrichment in *Enl*-T1 nephrons (**Figure 2c-f**). Furthermore, two clusters, namely C4 and C10 (hereafter referred to as T1-abnormal, or T1-ab), were markedly enriched in *Enl*-T1 nephrons (**Figure 2c-f**). Pseudotime trajectory analysis suggested a potential perturbation in nephron differentiation by *Enl*-T1, leading to two possible developmental paths from uncommitted NPCs (C0-NP1) (**Figure 2g**). One path resembles the differentiation trajectory typically observed in *Enl*-WT nephrons. However, most *Enl*-T1 cells may not progress past the committed NPCs (NP2) phase in this trajectory. An alternative path shows that NP1 cells might transition directly to T1-abnormal clusters (C4 and C10), with these clusters inferred to be poorly differentiated based on pseudotime scores (**Figure 2g**). Our results reveal the impact of *Enl*-T1 on the composition of nephron cells and their possible differentiation paths. We propose that *Enl*-T1 results in arrested nephrogenesis at early progenitor stages and the emergence of abnormal, undifferentiated progenitors (**Figure 2n**), a hypothesis that requires further validation through lineage tracing experiments.

### Transcriptional changes induced by mutant ENL in the developing kidney

To identify transcriptional changes induced by the mutant ENL in nephrons, we performed differential gene expression analysis between *Enl*-WT and *Enl*-T1 in cell populations that are reasonably abundant in the *Enl*-T1 nephron (NP1, NP2, and T1-ab). We confined our analysis of T1-abnormal cells to cluster 4, as cluster 10 was omitted due to a paucity of *Enl*-WT cells in this cluster (*n* < 50). We identified 1068 differentially expressed genes (DEGs, fold-change > 1.4), with a significant portion of DEGs shared among all five clusters examined (**Figure 2h**). Interestingly, while there were more downregulated than upregulated genes in T1 cells (**Figure S4a**), the upregulated DEGs exhibited a higher degree of changes in the percentage of cells expressing these genes (**Figure S4b**). Additionally, T1-upregulated (T1-UP) DEGs exhibited a higher degree of overlap across all five clusters (59 out of 182 DEGs) compared to downregulated (T1-DN) ones (122 out of 886 genes) (**Figure S4c, d**), suggesting that the impact of the mutation on the upregulated genes is more consistent.

Gene ontology (GO) analysis revealed that T1-UP DEGs were enriched in pathways associated with embryonic development, nephron development, and mitochondrial metabolism (**Figure 2i**). Among T1-UP DEGs involved in development, we observed a marked upregulation of numerous *Hox* genes across all five clusters (**Figure 2j**). In *Enl*-WT nephrons, *Hoxa* and *Hoxb* genes are ubiquitously expressed in the entire nephron, while *Hoxc* and *Hoxd* genes exhibit more cell-type specificity, with *Hoxc* expressed in NPCs and podocytes and *Hoxd* expressed in NPCs and DT (**Figure S4e**). *Hoxb* genes exhibited a relatively modest increase compared with the other three subfamilies, possibly due to high basal level in *Enl*-WT nephrons (**Figure S4f**). Among the *Hox* genes that are abnormally upregulated by *Enl*-T1, *Hox9, Hox10, Hox11* genes have been established as critical regulators of NPC self-renewal and precise execution of differentiation in genetic knockout studies^48–50^. This, together with our results, indicate that proper levels of *Hox* genes are crucial for kidney development. Consistent with scRNA-seq, RT-qPCR analyses of whole E15.5 kidneys revealed a pronounced upregulation of *Hox9/10/11* gene expression in *Enl*-T1 kidneys compared to *Enl*-WT counterparts (**Figure S4g**). Importantly, *ENL*-mutant human Wilms tumors exhibit elevated levels of certain *HOX* genes (e.g., *HOXA13*) compared to *ENL*-WT tumors^32^, supporting the clinical relevance of our findings and underscoring a potential role of *HOX* genes in Wilms pathogenesis. Intriguingly, genes involved in NPC commitment were also upregulated in *Enl*-T1 cells. For instance, *Wnt4*, which is typically expressed in the committed NPCs and functions as an inducer of MET^51,52^, had elevated expression in *Enl*-T1 NP1 (C0) and NP2 (C1/5) cells (**Figure 2j**). Furthermore, *Cdh6*, an epithelial marker gene usually expressed in the RV^42^, was aberrantly upregulated by *Enl*-T1 in all five clusters analyzed, including the NPCs (**Figure 2j**). These results suggest that *Enl*-T1 induces transcriptional changes involved in both self-renewal and cell fate commitment in nephron progenitors.

Quite interestingly, we also observed upregulation of genes related to mitochondria and metabolism in the nephron (**Figure 2i** and **Figure S4h**). Among these genes, the *Nduf*, *Atp*, and *Cox* gene families are linked to the respiratory electron transport chain (ETC)^53^. It has been shown that the ETC pathway becomes increasingly activated as nephron progenitors differentiate in the fetal kidney^54^, suggesting the possibility that *Enl*-T1 may promote premature commitment of NPCs in part through augmenting ETC activation. This hypothesis merits further exploration.

To determine whether the gene signatures are induced by *Enl*-T1 specifically in the nephron, we scored their expression across all major lineages using UCell, a tool for interrogating gene signatures in single-cell datasets^55^. Among *Enl*-T1-upregulated genes, the development-related signature was enhanced by *Enl*-T1 predominantly in the nephron and stroma, while the mitochondria-related signature was increased throughout the entire kidney (**Figure S4i**). Given that *Enl*-T1 expression was restricted to *Wt1*^+^ cells and their progeny in the nephron and stroma, these results suggest that the alteration of development-associated signatures is more likely a direct effect of *Enl*-T1 compared to the mitochondria-related signature.

We next assessed the clinical relevance of *Enl*-T1-induced gene signatures identified in our mouse model. We obtained RNA-seq datasets for Wilms tumor samples from the TARGET dataset^20^ and identified genes that are upregulated in ENL-mutant compared to ENL-wildtype tumors (Supplemental Table 3). We then performed gene set variation analysis (GSVA) to score both human and mouse ENL-mutant-UP signatures across all Wilms tumor samples and observed a positive correlation, suggesting their functional associations in human Wilms tumors (**Figure 2k**). Next, we assessed the expression of human ENL-mutant-UP signature in specific nephron cell types using our scRNA-seq datasets. *Enl*-T1 cells in the NP1, NP2, and T1-ab clusters exhibited higher expression levels of this signature compared to their wildtype counterparts, with the most significant increase occurring in T1-ab populations (**Figure 2l**). Collectively, these findings support the clinical relevance of our models in uncovering mutant ENL-induced transcriptional changes with implications for human Wilms tumors.

Next, we turned to genes that were downregulated in *Enl-*T1 nephrons. GO analysis revealed that T1-DN DEGs genes were associated with chromatin organization and transcription-related functions (**Figure 2m** and **Figure S4j**). Notably, *Dnmt1*, a methyltransferase essential for maintaining DNA methylation^56^, is known to be enriched in the nephrogenic zone of the developing kidney^57^. Studies have demonstrated that the loss of *Dnmt1* in nephron progenitors resulted in reduced self-renewal, leading to impaired renal differentiation and postnatal mortality in mice^57,58^. Additionally, key epigenetic regulators involved in histone methylation, such as *Ash2l* and *Ezh2*, which are expressed at higher levels in the *Six2*^+^ cap mesenchyme, were downregulated in *Enl*-T1 cells^59^. Ezh2 serves as the dominant H3K27 methyltransferase in *Six2*^+^ NPCs and is essential for their proper proliferation^60^. The reduced expression of these and other epigenetic regulators in *Enl*-T1 nephron precursors likely contributes to the observed nephrogenesis defects in *Enl*-T1 kidneys, a hypothesis that warrants further investigation. Like the *Enl-*T1-induced mitochondria-related signature, *Enl-*T1-downregulated genes found in the nephron showed a similar degree of decrease across different lineage compartments within the kidney (**Figure S4k**), suggesting a potential secondary effect of *Enl*-T1 expression.

### Chromatin accessibility dynamics and regulatory landscape during normal nephrogenesis

Dynamic control of chromatin accessibility plays a crucial role in cell differentiation by regulating the access of cell-type specific transcription factors (TFs) to their regulatory genomic regions^2,61^. To probe whether *Enl*-T1 affects the chromatin landscape in specific cell types, we performed single-nuclei Assay for Transposase Accessible Chromatin sequencing (snATAC-seq) on E15.5 *Enl*-WT and T1 kidneys. After applying a series of quality control metrics, we retained 17102 high-quality cells for subsequent analysis (**Figure S5a-e**). We first mapped the open chromatin landscape of *Enl*-WT kidneys. Based on the chromatin accessibly around the transcription start site (TSS) and gene body regions of well-known cell type-specific marker genes, we identified four major lineage compartments (**Figure S5f**) as in our scRNA-seq dataset. Within the nephron, cells were further grouped into several major cell types, including NPCs, intermediate cells, podocytes, PT, and DT/LOH (**Figure S5f, g**). Differential analysis identified 56,258 cell-type specific ATAC peaks across all eight major clusters within the kidney, with a majority exhibiting high specificity for a single cluster (**Figure S5h**). Motif enrichment analysis nominated top TFs that occupy these cell type-specific open regulatory elements, many of which also exhibited cell type-specific expression patterns (**Figure S5i, j**). Our results, together with recent scATAC-seq profiling of late-stage embryonic and postnatal kidneys^62,63^, define the cell type-specific chromatin and TF regulatory landscape during nephrogenesis in the mouse kidney.

To further understand the dynamic changes in chromatin accessibility that occur during cell fate transition, we defined major cell state transitions in *Enl*-WT nephrons based on differentiation trajectory using our snATAC-seq dataset (**Figure S6a-c**). We then identified differentially accessible regions (DARs) between descendant cell states along the trajectory (**Figure S6d**). Surprisingly, the transition from NP1 to NP2, signifying the commitment of NPCs, was characterized by a substantial gain in chromatin accessibility. Conversely, the transition from NP2 to podocytes showed a more pronounced loss in chromatin accessibility. The most substantial remodeling in chromatin accessibility occurred during the transition from NP2 to IM, coinciding with MET, a critical step for committed NPC to differentiate into various tubule structures^13–15^.

Motif enrichment analysis on the DARs identified candidate TFs that may regulate each step of cell fate transitions (**Figure S6e**). Upon examining the expression level and motif dynamic score using chromVAR^64^, we nominated 7 transcription factors (TFs), namely *Six2*, *Hoxc9*, *Wt1*, *Tcf21*, *Lhx1*, *Hnf1b* and *Hnf4a*, as key regulators orchestrating cell fate transitions during nephrogenesis with distinct dynamics (**Figure S6f, g**). For instance, *Six2* and *Hoxc9*, two NPC-specific TFs^40,62^, exhibited a gradual decrease in both their expression and binding site accessibility during the transition from the NP2 to subsequent stages. *Tcf21*, a key TF for glomerulogenesis^65^, was de novo activated during the NP2 to podocyte transition. *Lhx1* and *Hnf1b* were both highly expressed in IM cells, but they exhibited a mutually exclusive pattern of activity in subsequent lineages, with *Lhx1* activated in podocytes and DT and *Hnf1b* activated in PT. Thus, while an earlier study focusing on postnatal and adult kidneys highlighted the closing of open chromatin regions as the primary event during nephron differentiation^62^, our findings uncover novel aspects of chromatin dynamics and key regulatory TFs during cell fate transitions in the developing mouse kidney.

### *Enl*-mutant kidneys exhibit an altered open chromatin landscape during early nephrogenesis

Having established the open chromatin landscape of the *Enl*-WT embryonic kidneys, we investigated how this landscape is impacted by *Enl*-T1. Through the integration of snATAC-seq datasets obtained from *Enl*-WT and T1 kidneys, we found that *Enl*-T1 kidneys exhibited a reduction in specific nephron clusters and an increase in certain stromal clusters (**Figure 3a, b**). Through label transfer^66^, we used scRNA-seq annotations as a reference to assign cell type identity to nephron cells in snATAC-seq datasets (**Figure 3c**). This analysis revealed several intriguing observations. First, while *Enl*-WT nephron cells displayed a variety of chromatin states corresponding to cell types found in nephrogenesis, *Enl*-T1 cells exhibited a marked reduction in chromatin state diversity. In particular, *Enl*-T1 cells showed a substantial loss of chromatin states associated with differentiated nephron structures (e.g., PT, DT, Podo) (**Figure 3d, e**) and a more uniform chromatin state of clusters representing undifferentiated cells. Second, while *Enl*-T1 NP1 cells clustered with *Enl*-WT counterparts based on gene expression (**Figure 2c, d**), they exhibited a shift towards NP2 cells (C1/5/7) in the snATAC-seq dataset (**Figure 3c**). This observation was further supported by a lower number of DARs between NP1 and NP2 in *Enl*-T1 (NP2 vs. NP1: 2153 gained and 267 lost) compared to the DARs in *Enl*-WT nephrons (NP2 vs. NP1: 11020 gained and 547 lost). These findings suggest that *Enl*-T1 NP1 cells may have substantial chromatin alterations before overt transcriptional changes occur. Furthermore, although scRNA-seq analysis indicated that NP2 (C1/5/7) and T1 abnormal clusters (C4/10) in *Enl*-T1 nephrons follow distinct differentiation paths originating from NP1 (**Figure 2g**), a higher degree of similarity in chromatin accessibility was observed between these clusters (**Figure 3c**). These results reveal dynamic changes in chromatin accessibility induced by the *Enl* mutation, shedding light on aspects that would otherwise remain masked in scRNA-seq analysis alone.

**Figure 3.**
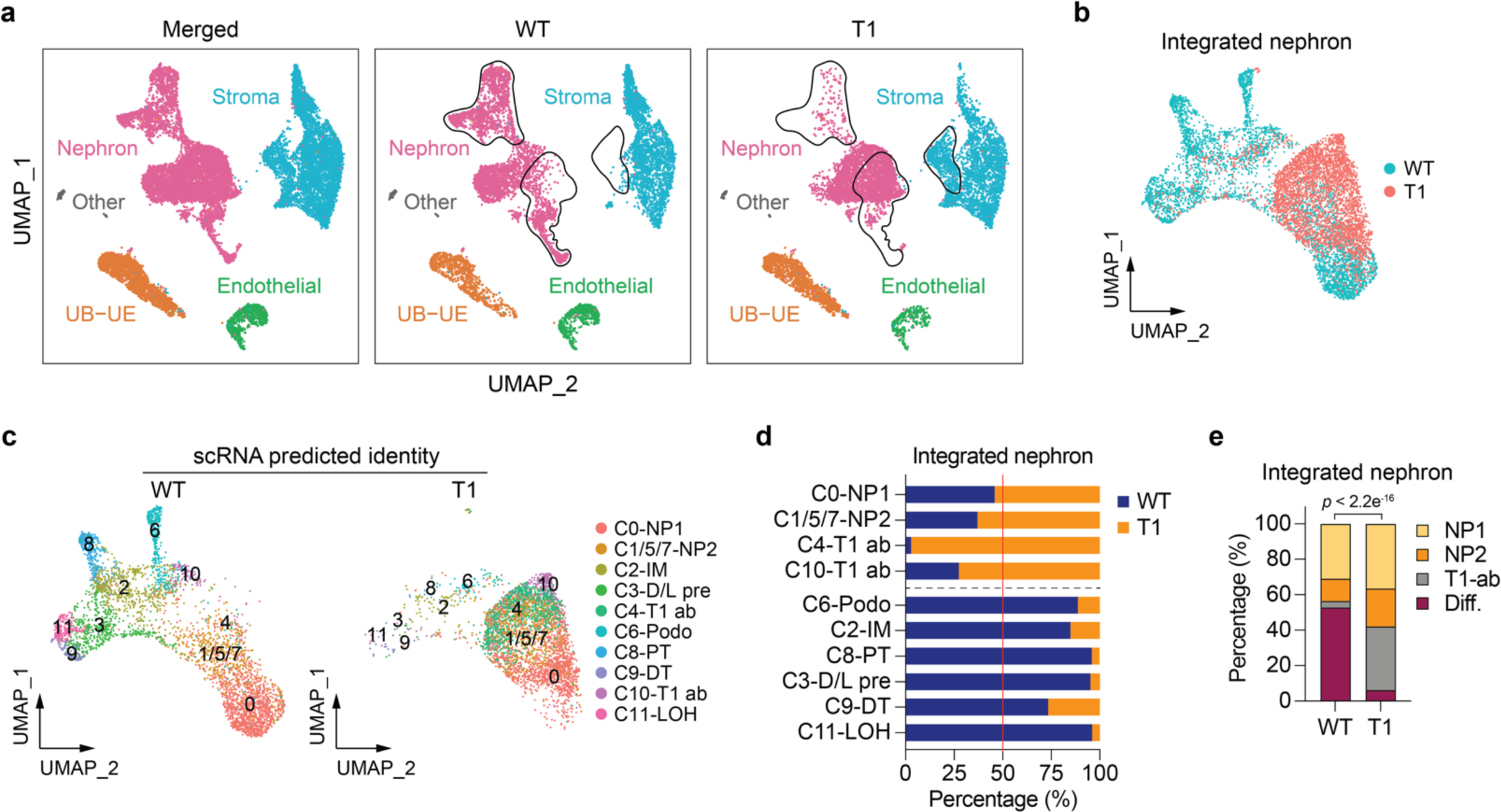
snATAC-seq reveals changes in the open chromatin landscape in *Enl*-mutant kidneys during nephrogenesis. **a**, UMAP embedding of snATAC-seq data labeled with four main embryonic kidney lineages. The dramatically altered clusters between *Enl*-WT and T1 kidneys are highlighted with black circles. **b**, UMAP embedding of integrated snATAC-seq cells from *Enl*-WT and T1 nephrons, colored by sample. **c**, UMAP embedding of integrated snATAC-seq cells from *Enl*-WT (left) and T1 (right) nephrons, respectively. Cells are colored and labeled by the cell types predicted by corresponding scRNA-seq data. **d,** Stacked bar plot showing the percentage of scRNA-seq predicted cell types between *Enl*-WT and T1 nephrons. **e**, Stacked bar plot showing the percentage of scRNA-seq predicted cell types within *Enl*-WT and T1 nephrons. Chi-Square test *p*-value is shown. Diff., differentiation, including RV, Podo, PT, D/L pre, DT, and LOH cell types.

### Mutant ENL promotes premature commitment of nephron progenitors while simultaneously restricting their differentiation through misregulation of specific TF regulons

To understand the regulatory mechanism underlying altered chromatin states in *Enl*-T1 uncommitted NPCs, we compared chromatin accessibility between WT and T1 NP1 cells. We identified 8958 gained and 5616 lost DARs in *Enl*-T1 cells (**Figure 4a**), the majority of which were in distal intergenic regions or gene bodies, likely representing enhancers (**Figure S7a**). GREAT analysis (**Figure 4b** and **Figure S7b**) revealed that T1-gained, but not T1-lost, DARs were strongly associated with functional terms related to nephrogenesis, such as metanephric nephron morphogenesis and renal vesicle morphogenesis, indicating that *Enl*-T1 NP1 cells are more primed to commitment and differentiation at the chromatin level. Motif enrichment coupled with gene expression analysis identified several *Hox* TFs (*Hoxa9*, *Hoxc9*, *Hoxa11*, and *Hoxd11*) as top regulators of gained DARs (**Figure 4c**). Notably, *Hoxc9* has been identified as a key TF driving the NP1 to NP2 transition during normal nephrogenesis (**Figure S6f**). In contrast, expression levels and motif accessibility of several TFs important for NPC self-renewal, such as *Six2*^40^ and *Wt1*^36^, were decreased in *Enl*-T1 NP1 cells, with *Six2* standing out as the most significant hit (**Figure 4c**). These results suggest that mutant ENL may disrupt the balance of self-renewal and commitment of NPCs through misregulation of key TF regulons and prime these cells for differentiation at the chromatin level. Furthermore, we found that 20% of *Enl*-T1 gained DARs in NP1 cells coincided with open chromatin regions associated with the NP1 to NP2 transition during normal nephrogenesis (**Figure S6d** and **Figure S7c**), suggesting that *Enl*-T1 promotes premature commitment of NP1 cells through the acquisition of normal development-associated as well as de novo open chromatin regions.

**Figure 4.**
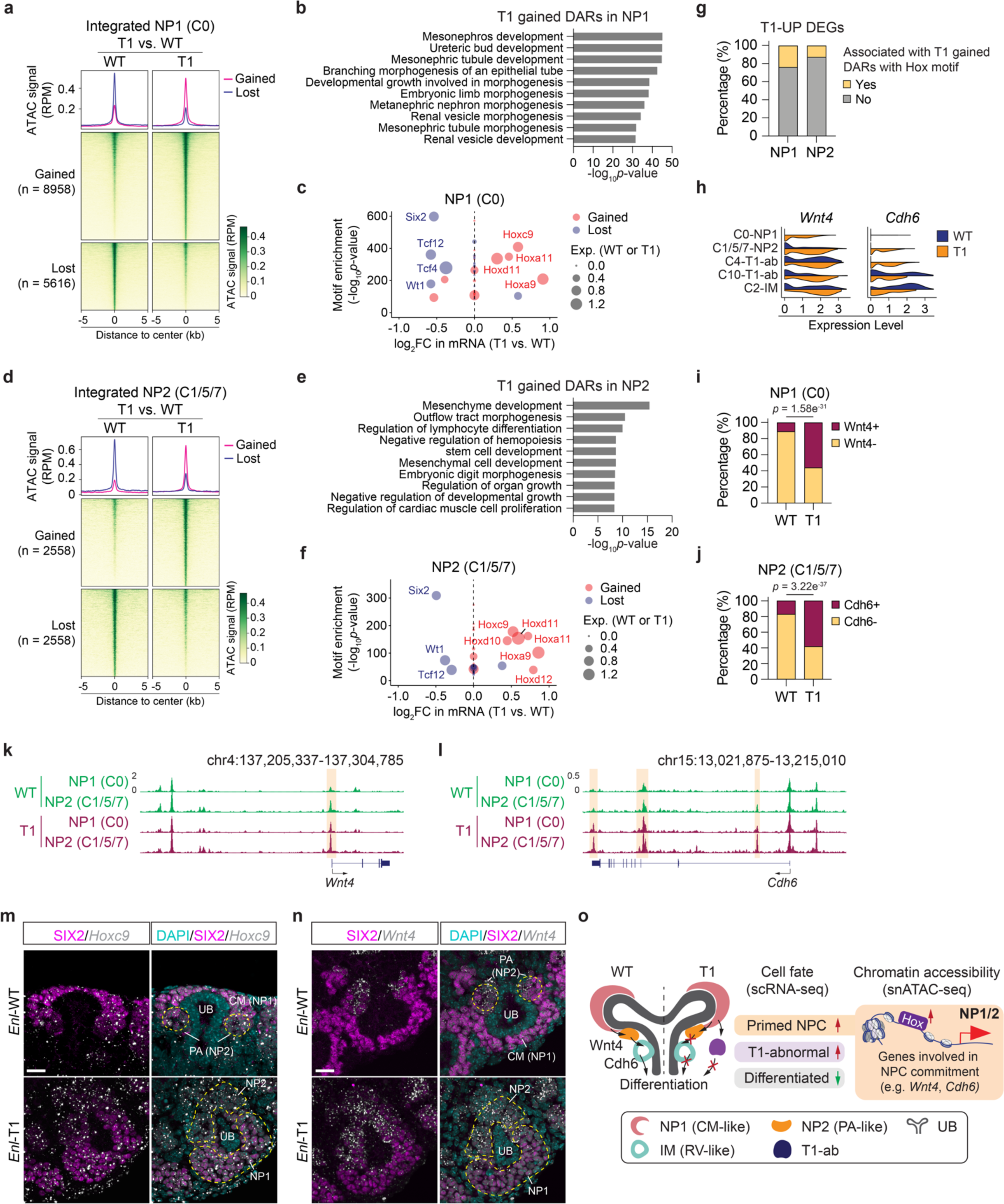
Mutant ENL promotes premature commitment of nephron progenitors while simultaneously restricting their differentiation through misregulation of specific TF regulons. **a**, **d**, snATAC data in NP1 (**a**) and NP2 (**d**) are plotted as average occupancies (top) and heatmap (bottom) across the differentially accessible regions (DARs) between *Enl*-WT and T1 cells. The ATAC signal is normalized by reads per million (RPM). All regions are defined as gained (up-regulated in *Enl*-T1, red) and lost (down-regulated in *Enl*-T1, blue), and the corresponding numbers are shown on the left. See Supplementary Table 7. **b**, **e**, GREAT analysis of T1-gained DARs in NP1 (**b**) and NP2 (**e**). **c**, **f**, Dot plot showing the top 20 most significant TFs identified from the motif enrichment analysis for T1 gained or lost DARs in NP1 (**c**) and NP2 (**f**). Fold changes between *Enl*-WT and T1 for each TF are shown in the x-axis. Scaled *p*-values of motif enrichment for each TF are shown in y-axis. The TF candidates identified from T1 gained or lost DARs are colored in red or blue, respectively. The expression level in *Enl*-WT (blue) or T1 (red) cells for each TF is represented by the size of the circle. FC, fold-change. Exp., expression. **g**, Stacked bar plots indicating the percentage of T1-UP DEGs associated with Hox motif enriched T1 gained DAR. **h**, Violin plots showing the expression level of *Wnt4* and *Cdh6* for NP1, NP2, T1-ab, and IM cell types in *Enl*-WT or T1 nephrons. **i**, Stacked bar plots indicating the percentage of NP1 cells with or without *Wnt4* expression in *Enl*-WT or T1 nephrons. Fisher’s exact test *p*-value is shown. **j**, Stacked bar plots indicating the percentage of NP2 cells with or without *Cdh6* expression in *Enl*-WT or T1 nephrons. Fisher’s exact test *p*-value is shown. **k**, **l**, The genome browser view of ATAC signals at *Wnt4* (**k)** and *Cdh6* (**l**) gene loci in indicated cell types from *Enl*-WT (top) or T1 (bottom) nephrons. **m**, Representative images of SIX2/*Hoxc9* mRNA co-staining in E15.5 kidneys. CM, cap mesenchyme; UB, ureteric bud; PA, pretubular aggregate. Scale bar = 20 µm. **n**, Representative images of SIX2/*Wnt4* mRNA co-staining in E15.5 kidneys. CM, cap mesenchyme; UB, ureteric bud; PA, peritubular aggregate. Scale bar = 20 µm. **o**, Schematic illustrating impaired nephrogenesis phenotypes and potential molecular mechanism we identified in the *Enl*-T1 NP1 and NP2.

Despite the premature commitment of *Enl*-T1 NP1 cells to a NP2-like state, these cells failed to give rise to differentiated kidney structures. To explore the underlying mechanisms, we compared the chromatin accessibility between *Enl*-WT and *Enl*-T1 NP2 cells. As in the case of T1-induced DARs in NP1 cells, DARs identified in NP2 cells were primarily located in distal regulatory regions (**Figure 4d** and **Figure S7d**). However, GREAT analysis revealed that these DARs were not directly related to nephrogenesis but instead, they exhibited strong enrichment in mesenchyme development pathways (**Figure 4e** and **Figure S7e**). Given that normal NP2 represents primed NPCs in the pretubular aggregates that undergo MET, a prerequisite for nephron differentiation, our results suggest that *Enl*-T1 NP2 cells may exhibit a defect in differentiation due to aberrant maintenance of a mesenchymal chromatin state. Furthermore, an integrative analysis of TF motif enrichment and expression revealed similar regulators in *Enl*-T1 NP2 cells as those identified in NP1 cells (**Figure 4f**), underscoring the sustained impact of the mutant on the chromatin state of self-renewing and committed nephron progenitors during early nephrogenesis. Interestingly, our scATAC-seq on *Enl*-WT embryonic kidneys revealed that both the expression levels and activity of *Hoxc9* gradually decrease during the transition from NP2 to IM (**Figure S6f, g**), a precursor stage for podocyte and tubule differentiation (**Figure S3h**). Therefore, we speculated that the persistently elevated expression and activity of *Hoxc9* and possibly other *Hox* TFs contribute to the impeded differentiation of *Enl*-T1 NP2 cells. To address this hypothesis, we performed a regulon analysis to identify putative target genes of *Hox* TFs in NP1 and NP2 cells. The results showed that > 80% of *Enl*-T1-gained DARs contained at least one motif sequence of identified *Hox* TFs (**Figure S7f**). We then identified genes associated with *Hox* motif-containing DARs and examined their overlap with T1-upregulated DEGs. We found that 24% of T1-upregulated DEGs in NP1 cells and 13% in NP2 cells have the potential to be activated by *Hox* TFs (**Figure 4g** and **Figure S7g**, **h**). Notably, among these targets, *Wnt4*, essential for NPC commitment and MET initiation^51,52^, and *Cdh6*, an epithelial marker in RV^42^, showed aberrant activation in *Enl*-T1 NP1 and NP2 cells, respectively (**Figure 4h**). Over 55% of *Enl*-T1 NP1 cells expressed *Wnt4*, compared to only ∼10% of *Enl*-WT cells (**Figure 4i**). Similarly, the percentage of *Cdh6*^+^ cells in NP2 cells was increased from 17.1% in *Enl*-WT to 58.4% in *Enl*-T1 (**Figure 4j**). Furthermore, we observed an increase in chromatin accessibility at the *Wnt4* and *Cdh6* gene loci, which likely contributes to their elevated expression (**Figure 4k, l**).

To validate transcriptional changes identified by scRNA-seq, we performed RNA *in situ* hybridization (ISH) and/or immunostaining for several key genes in E15.5 kidneys. First, we focused on *Hoxc9*, a top transcription factor candidate predicted to mediate increased chromatin accessibility in *Enl*-T1 *Six2*^+^ NPCs (**Figure 4b** and **4e**). *Hoxc9* mRNA ISH coupled with SIX2 protein co-staining showed that *Hoxc9* was highly expressed in SIX2^+^ NPCs in *Enl*-WT kidneys (**Figure 4m**), consistent with our scRNA-seq data (**Figure S4e**). Notably, *Hoxc9* ISH signals were elevated in SIX2^+^ NPCs in *Enl*-T1 kidneys (**Figure 4m**). We then aimed to confirm the upregulation of *Wnt4* and *Cdh6* in *Enl*-T1 NP1 and NP2 cells. Imaging data showed that in *Enl*-WT kidneys, *Wnt4* was present in SIX2^low^ cells within the PA structure, likely corresponding to NP2 cells, and absent in SIX2^high^ cells in the CM structure, likely corresponding to NP2 cells (**Figure 4n** and **Figure S7i**). This pattern aligns well with our scRNA-seq data (**Figure 4h**) and previous findings^67^. We observed increased *Wnt4* ISH signals in *Enl*-T1 SIX2^+^ NP1 and NP2 cells relatively to *Enl*-WT (**Figure 4n** and **Figure S7i**). During normal kidney development, *Cdh6* expression begins in the intermediate (IM) cells within the renal vesicle and subsequently appears in differentiated cells (**Figure 4h**). CDH6 immunostaining confirmed that in *Enl*-WT kidneys, CDH6 was not detected in NPCs located in the CM and PA structures, but it was present in cells within the RV structure, which likely represent the IM cells identified in our scRNA-seq analysis. In contrast, in *Enl*-T1 kidneys, *Cdh6* was aberrantly expressed in NP2 cells, as revealed by scRNA-seq (**Figure 4h**), and CDH6 protein was detected within the PA structure (**Figure S7j**). These *in situ* validation results lend further support to findings from scRNA-seq. We propose that the *Hox*-mediated increase in chromatin accessibility and gene expression of key developmental genes, such as *Wnt4* and *Chd6*, may underlie the aberrant commitment of *Enl*-T1 NPCs (**Figure 4o**).

### An abnormal progenitor state losing nephron chromatin identity appears in *Enl*-mutant kidneys

Next, we aimed to characterize the abnormal cell cluster (C4) gained in *Enl*-T1 nephron. Trajectory analysis on the scATAC-seq dataset placed NP2 and T1-ab cells downstream of NP1 cells, and NP2 and T1-ab cells exhibited similar differentiation scores (**Figure 5a** and **Figure S8a**). These results might suggest that NP2 and T1-ab cells represent subsets of primed NPCs. Moreover, trajectory analysis based on scRNA-seq dataset points to a potential divergence from the self-renewing NP1 to either NP2 or T1-ab cells, highlighting differences in the cellular state between these two populations (**Figure 2h** and **Figure S8b, c**). To explore the mechanism underlying the emergence of T1-abnormal cells, we identified DARs between the predominant T1-ab cluster (C4) and NP1 (C0) or NP2 (C1/5/7) in *Enl*-T1 nephrons. The analysis revealed substantial differences in open chromatin landscapes between T1-ab and NP1, with 1109 gained and 2518 lost DARs in T1-ab cells (**Figure 5b**). In contrast, when comparing T1-ab and NP2, we observed a decrease in ATAC signals in only 629 chromatin regions (**Figure 5c**). These results suggest a more similar chromatin state between T1-ab and NP2 cells compared to the differences observed between T1-ab and NP1 cells. GREAT analysis on these DARs showed that T1-ab cells lost the potential for nephrogenesis compared with NP1 (**Figure 5d**) and NP2 cells (**Figure 5e**). Specifically, when compared to NP2 cells, DARs lost in T1-ab cells were enriched for GO terms related to renal vesicle morphogenesis and mesenchymal to epithelial transition, indicating a more pronounced defect in their differentiation into renal vesicles (**Figure 5e**). Intriguingly, T1-ab cells gained the potential at the chromatin level to differentiate into other organs (**Figure 5f**). These results suggest that the mutant ENL induces an abnormal progenitor state characterized by a loss of open chromatin regions encoding nephron identity or differentiation potential.

**Figure 5.**
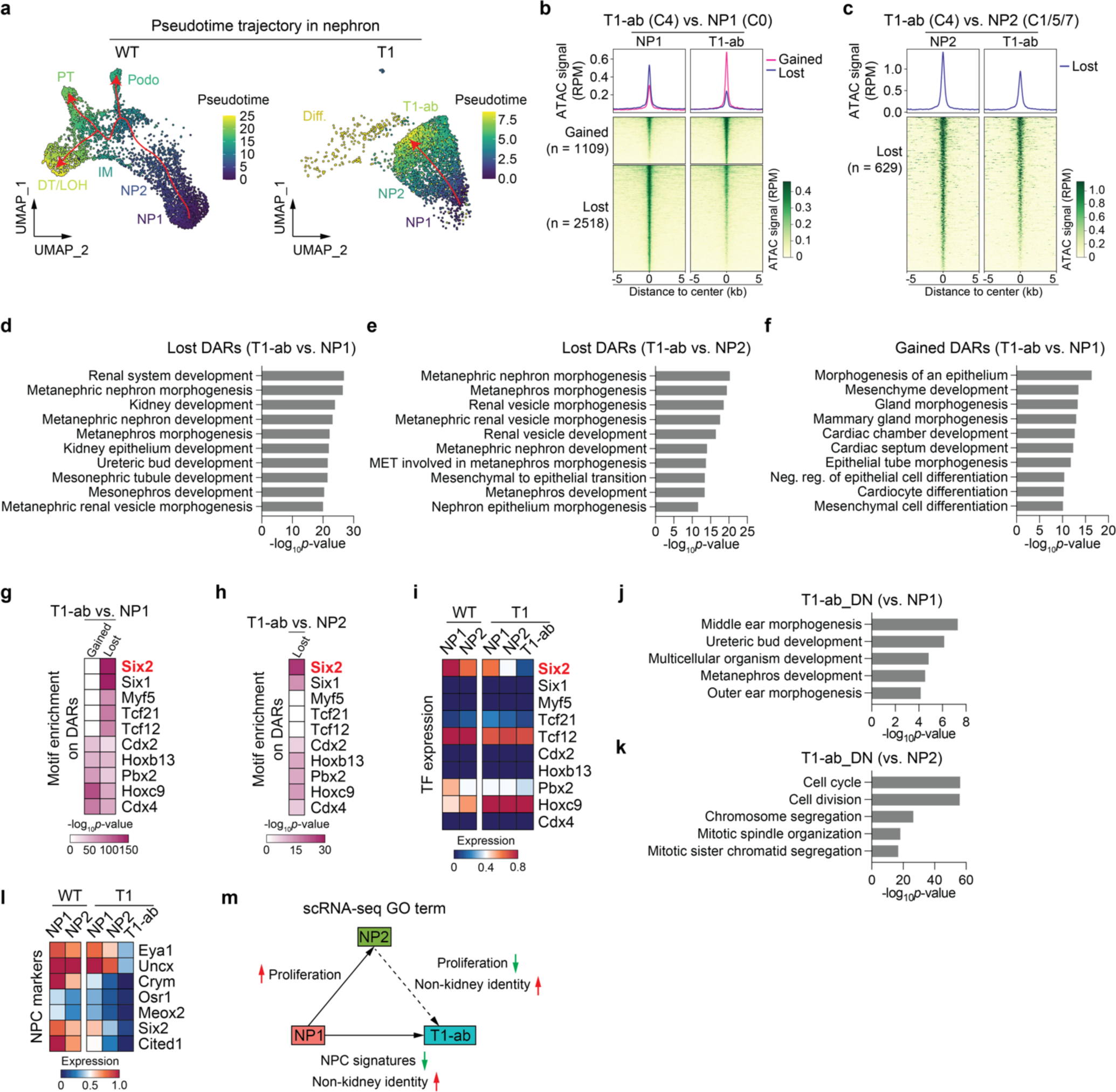
An abnormal progenitor state losing nephron chromatin identity emerges in *Enl*-mutant kidney. **a**, UMAP embedding of integrated *Enl*-WT (left) or T1 (right) snATAC-seq nephron differentiation trajectory. Cells are colored by pseudotime. Trajectories are depicted by red arrows. **b**, snATAC data in *Enl*-T1 for NP1 (left) and T1-ab (right) are plotted as average occupancies (top) and heatmap (bottom) across the DARs between NP1 and T1-ab cells. The ATAC signal is normalized by RPM. All regions are defined as gained (up-regulated in T1-ab, red) and lost (down-regulated in T1-ab, blue), and the corresponding numbers are shown on the left. See Supplementary Tables 9. **c**, snATAC data in *Enl*-T1 for NP2 (left) and T1-ab (right) are plotted as average occupancies (top) and heatmap (bottom) across the T1-ab lost DARs. The ATAC signal is normalized by RPM. The corresponding DAR numbers are shown on the left. See Supplementary Tables 9. **d**-**f**, Bar plots showing GREAT analysis for T1-ab vs. NP1 lost **(d)**, T1-ab vs. NP2 lost **(e)**, and T1-ab vs. NP1 gained **(f)** DARs in *Enl*-T1 cells. **g**, **h**, Heatmap showing the motif enrichment *p*-value of the top TF candidates identified from the motif analysis for the DARs indicated in (**b)** and (**c**). *p*-values are scaled by -log10. **i**, Heatmap showing the expression level of TFs identified in (**g**) and (**h**) in *Enl*-WT and *Enl*-T1 NP1, NP2, and T1-ab scRNA-seq cells. **j**, **k**, Bar plots showing GO term analysis for T1-ab vs. NP1 DN DEGs **(j)** and T1-ab vs. NP2 DN DEGs **(j)** in the *Enl*-T1 nephron. **l**, Heatmap showing the expression levels of NPC markers in indicated cell types of *Enl*-T1 nephron. **m**, Schematic summarizing the GO term analyses performed in (**j**), (**k**), and Figure S**8g-i**.

To identify TFs that regulate chromatin changes in T1-ab cells, we performed motif enrichment analyses on identified DARs. There was a strong enrichment of motifs for several TFs involved in NPC self-renewal and maintenance, such as *Six2*^40^ and *Tcf21*^68^, in T1-ab lost DARs when compared with NP1 cells (**Figure 5g**). Similar results were observed in T1-ab vs. NP2 comparison, albeit to a lesser extent, which may be attributed to a lower number of DARs (**Figure 5h**). Among these TFs, *Six2* exhibited the most significant gene expression alteration among NP1, NP2, and T1-ab cells, with T1-ab cells showing the lowest expression levels (**Figure 5i** and **Figure S8d**). As the *Six2* gene locus displayed minimal changes in ATAC signals across these three populations (**Figure S8e**), downregulation of its upstream regulators or changes in other chromatin features may underlie the observed expression changes. Nevertheless, the loss of *Six2* binding regions observed in T1-ab cells likely contributes, at least in part, to the reduced potential of T1-ab to specify the kidney lineage.

On the other hand, several TFs, including *Hoxc9*, *Hoxb13*, *Cdx2*, *Pbx2*, and *Cdx4*, were identified as top candidates that potentially regulate T1-ab gained DARs when compared with NP1 cells (**Figure 5g**), despite their similar expression levels in T1-ab and NP1 cells (**Figure 5i**). Intriguingly, motifs for these same TFs were also enriched, albeit to a lesser extent, in T1-ab lost DARs (vs. NP1) (**Figure 5g**). These TFs may cooperate with distinct partners to induce the opening or closing of different chromatin regions. Among these TFs, *Hoxc9* ranked as the top TF candidate occupying T1-ab gained DARs. We have demonstrated that *Hoxc9* is a key TF driving NPC commitment (NP1 to NP2 transition) during normal nephrogenesis (**Figure S6e, g**). As both T1-ab and T1-NP1 cells exhibited aberrantly high levels of *Hoxc9*, we speculated that *Hoxc9* may promote the opening of additional chromatin regions in T1-ab cells that regulate other lineage specifications, a hypothesis requiring further investigation.

Having determined the open chromatin landscape in T1-ab cells, we next investigated the transcriptional changes underlying their aberrant cell state. We identified DEGs across T1-ab, NP1, and NP2 cells in *Enl*-T1 nephrons (**Figure S8f**) and performed GO term analyses for these DEGs. Genes that were downregulated in T1-ab cells compared to NP1 were enriched in pathways related to kidney development (**Figure 5j**). Notably, the expression levels of several well-known NPC markers were markedly decreased in T1-ab cells, consistent with these cells losing the NPC state (**Figure 5l**). Additionally, compared to NP2, T1-ab cells exhibited a defect in proliferation, a cellular process required during normal NPC commitment^69^ (**Figure 5k** and **Figure S8g**). Furthermore, genes upregulated in T1-ab cells compared to either NP1 or NP2 are associated with the development of other tissues/organs or in metabolism (**Figure 5m** and **Figure S8h, i**). Integrated analysis of scRNA-seq and scATAC-seq datasets revealed that only a subset of DEGs was linked to changes in chromatin accessibility (**Figure S8j**), suggesting additional mechanisms involved in transcriptional regulation. Taken together, our chromatin and transcriptional analyses indicate that T1-abnormal cells are arrested in an aberrant progenitor state characterized by a loss of kidney lineage identity and a gain of potential to develop into other lineages (**Figure 5m**).

### Mutant ENL alters the open chromatin landscape of *Foxd1*^+^ stromal progenitors

The kidney stroma plays a crucial role in renal morphogenesis through interactions with the nephron and ureteric bud^17–19,45^. However, the precise origins, developmental hierarchy, and regulatory mechanisms of the stroma remain poorly understood. Our scRNA-seq analyses have shown that *Enl*-T1-induced development-related gene signatures were also found in the stroma (**Figure S4i**). Moreover, *Enl*-T1 stroma exhibited substantial changes in the open chromatin landscape compared to *Enl*-WT stroma (**Figure 3a**). Therefore, we sought to investigate how alterations in the stroma may contribute to the observed nephrogenesis defects in *Enl*-T1 kidneys.

Unbiased clustering of stroma cells extracted from our scRNA-seq datasets resulted in 10 clusters (**Figure 6a, b**) representing 6 distinct cell types, including *Foxd1*^+^ stroma progenitor (SP) (*Foxd1*^+^, *Dlk*^+^), proliferating cortical stroma (CS) (*Top2a*^+^), CS (*Clca3a1*^+^), medullary stroma (MS) (*Alx1*^+^, *Wnt4*^+^), ureteric stroma (US) (*Myh11*^+^), and renin/mesangial cells (Ren/Mes) (*Ren1*^+^, *Akr1b7*^+^) (**Figure S9a**). The three major stroma subdomains, CS, MS, and US, which exhibit a dorsoventral distribution pattern (CS, dorsal; MS and US, ventral) in the developing kidney^46^, were well separated in our scRNA-seq datasets (**Figure 6a, b**), indicating distinct gene expression and functions of these subdomains. Trajectory analysis based on scRNA-seq datasets on *Enl*-WT stroma (**Figure S9b**) suggested that *Foxd1*^+^ SP (C0/C8) is the origin of CS/MS populations, which constitute most of the kidney interstitium^18^. In contrast, the US cells (C5) represented a distinct population derived from *Tbx18*^+^ SP, consistent with previous lineage tracing studies^70^. Mutant ENL did not significantly alter the developmental trajectories of stroma cells.

**Figure 6.**
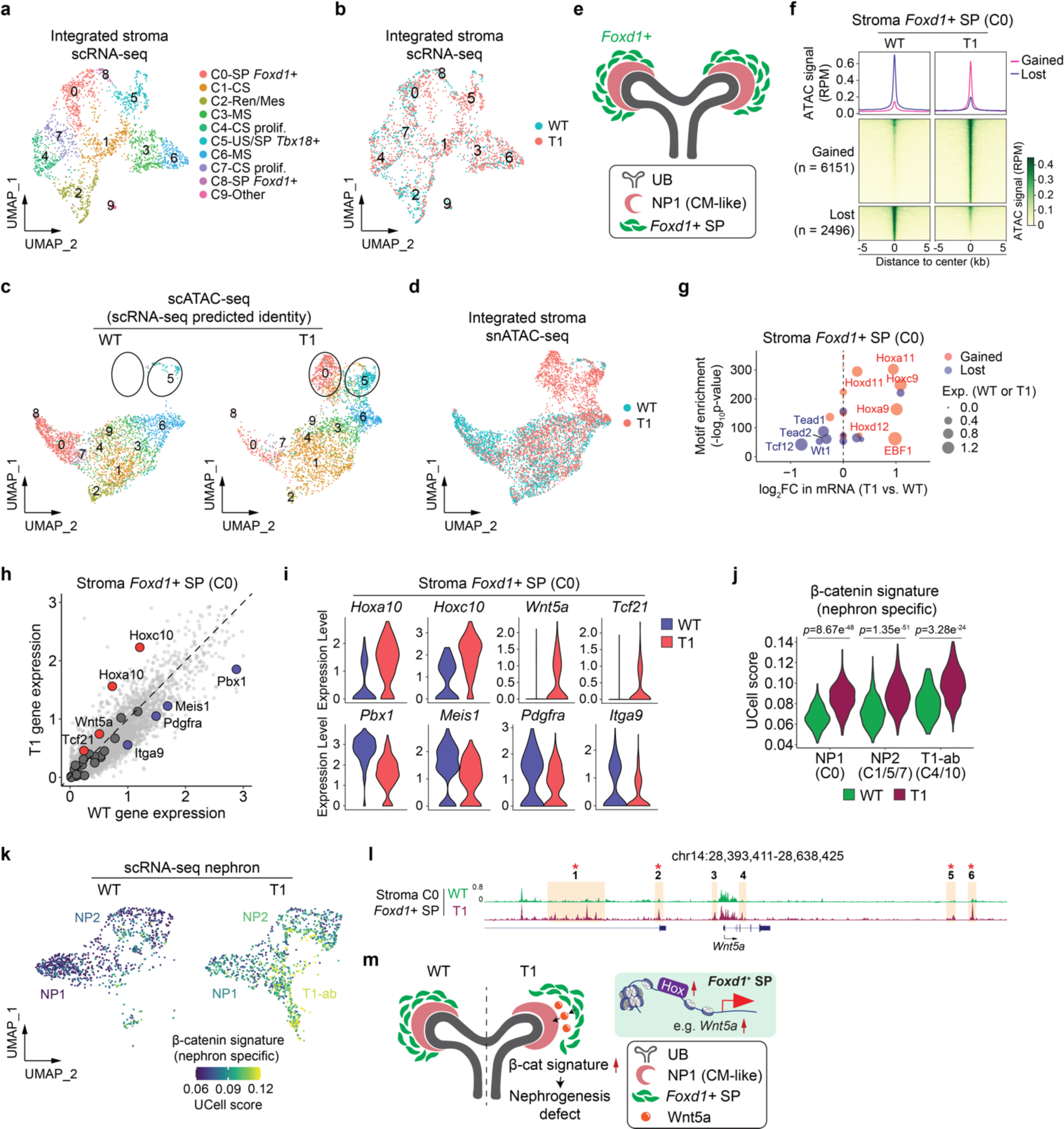
*Enl*-mutant *Foxd1*^+^ stromal progenitors exhibit altered chromatin accessibility and might affect stroma-nephron interactions by aberrantly activating paracrine Wnt signaling. **a**, UMAP embedding of integrated scRNA-seq cells from *Enl*-WT and T1 stroma. Cells are colored and labeled by annotated cell types. SP, stroma progenitor; CS, cortical stroma; Ren/Mes, renin/mesangial cells; MS, medullary stroma; prolif., proliferation; US, ureteric stroma. **b**, UMAP embedding of integrated scRNA-seq cells from *Enl*-WT and T1 stroma. Cells were colored by sample. **c**, UMAP embedding of integrated snATAC-seq cells from *Enl*-WT (left) or T1 (right) stroma. Cells are colored and labeled by the cell types predicted by corresponding scRNA-seq data. The dramatically altered clusters (C0 and C5) between *Enl*-WT and T1 stroma are highlighted with black circles. **d**, UMAP embedding of integrated snATAC-seq cells from *Enl*-WT and T1 stroma. Cells were colored by sample. **e**, Schematic illustrating the spatial arrangement of *Foxd1^+^* SP and CM-like population NP1 within the kidney. **f**, snATAC data in *Enl*-WT and T1 stroma C0 are plotted as average occupancies (top) and heatmap (bottom) across the differentially accessible regions (DARs) between *Enl*-WT and T1 cells. The ATAC signal is normalized by reads per million (RPM). All regions are defined as gained (up-regulated in *Enl*-T1, red) and lost (down-regulated in *Enl*-T1, blue), and the corresponding numbers are shown on the left. See Supplementary Table 12. **g**, Dot plot showing the top 20 most significant TFs identified from the motif enrichment analysis for T1 gained or lost DARs in stroma C0. Fold changes between *Enl*-WT and T1 for each TF are shown in the x-axis. Scaled *p*-value of motif enrichment for each TF is shown in y-axis. The TF candidates identified from T1 gained or lost DARs are colored in red or blue, respectively. The expression level in *Enl*-WT (blue) or T1 (red) cells for each TF is represented by the size of the circle. FC, fold-change. Exp., expression. **h**, Scatter plot showing the expression level of the whole mouse genome in *Enl*-WT and T1 stroma C0. 28 stroma-nephron interaction-related genes implicated or predicted previously are highlighted. Among them, T1-UP DEGs are colored in red, T1-DN DEGs are colored in blue, and no-change genes are highlighted in grey. See Supplementary Table 13. **i**, The expression level of indicated DEGs highlighted in (**h**) in *Enl*-WT and T1 stroma C0. **j**, The UCell score evaluated by nephron specific β-catenin activation signature for the integrated cell types of NP1, NP2, and T1-ab within *Enl*-WT and T1 nephrons. Wilcoxon rank-sum test *p*-values are shown. **k**, UMAP embedding of scRNA-seq NP1, NP2, and T1-ab cells showing the nephron specific β-catenin signature in *Enl*-WT or T1. Cells are colored by the UCell score of the signature. **l**, ATAC signals at *Wnt5a* gene locus in stroma C0 from *Enl*-WT or T1. T1-gained DARs are highlighted and numbered, in which DAR containing Hox TF motif is indicated with star. **m**, Schematic illustrating the model that the nephrogenesis defects in *Enl*-T1 kidney may be partially attributed to the abnormal hyper-activation of nephrogenic β-catenin due to Hox-driven up-regulation of *Wnt5a* in *Foxd1*^+^ SP.

To gain a comprehensive understanding of chromatin changes induced by mutant ENL in the stroma, we assigned cell types to snATAC-seq datasets based on the corresponding annotations from the scRNA-seq dataset. Several notable chromatin changes in *Enl*-T1 stroma were identified, including an embedding shift of the *Foxd1*^+^ SP cells (C0) and an expansion of the US cluster (C5) in the UMAP (**Figure 6c, d**). Histological studies on the mouse embryonic kidney^71^ showed that *Foxd1*^+^ SP, but not *Tbx18*^+^ SP, constitutes the stroma population surrounding the uncommitted NPC-containing cap mesenchyme (CM) (**Figure 6e**). This observation, coupled with reported interactions between *Foxd1*^+^ SP and CM, led us to speculate that abnormal *Foxd1*^+^ SP cells in *Enl*-T1 kidneys may impact nephrogenesis by influencing stroma-nephron interactions. Re-clustering of extracted *Foxd1*^+^ SP cells (C0) from the integrated stroma snATAC-seq datasets confirmed that, while a small portion of *Enl*-T1 *Foxd1*^+^ SP cells clustered with *Enl*-WT counterpart, the majority exhibited a distinct open chromatin landscape (**Figure S9c**). Interestingly, we did not observe a significant separation of these cells from *Enl*-WT counterparts based on scRNA-seq data (**Figure 6b**), suggesting that chromatin changes in these cells precede overt transcriptional alterations. Differential analysis of *Foxd1*^+^ SP cells (C0) between T1 and WT identified 6151 T1-gained and 2496 T1-lost DARs (**Figure 6f**). Interestingly, GREAT analysis of the gained DARs revealed an enrichment for GO terms associated with nephron and UB development (**Figure S9d**). Several well-known NPC and UB marker genes, such as *Cited1*^16^ and *Ret*^72^, exhibited increased chromatin accessibility (**Figure S9e**). These results suggest that *Enl*-T1 *Foxd1*^+^ SP cells may acquire lineage plasticity at the chromatin level. Furthermore, the DARs lost in *Enl*-T1 cells were associated with epithelial-to-mesenchymal transition (EMT) (**Figure S9d**), raising the possibility that these cells might have a compromised ability to maintain their mesenchymal state. Integrating motif and gene expression analysis, we identified *Hox9/11* TFs as top candidates that bind to T1-gained DARs and have higher expression levels in *Enl*-T1 *Foxd1*^+^ SP cells (**Figure 6g**). Previous research has demonstrated the role of stroma-derived *Hox10* paralogs in renal morphogenesis by affecting the reciprocal interaction between *Foxd1*^+^ SP cells and nephrogenic mesenchyme^73^. While we found that *Hox10* genes, especially *Hoxa10* and *Hoxc10*, were highly expressed in *Enl*-T1 *Foxd1*^+^ SP cells, the expression levels of *Hox9/11* subfamilies were much more elevated by mutant ENL (**Figure S9f**). In line with these results, *Hoxc9* RNA ISH coupled with SIX2 protein co-staining revealed increased *Hoxc9* expression in the *Six2*-negative stroma cells at the periphery of the *Six2*^+^ cap mesenchyme (**Figure 4m**), which likely correspond to the *Foxd1*^+^ stroma progenitors^71^. Altogether, these results suggest a potential role of *Hox9/11* in modulating the chromatin state of *Enl*-T1 *Foxd1*^+^ SP cells.

### *Enl*-mutant *Foxd1*^+^ stromal progenitors might affect stroma-nephron interactions by aberrantly activating paracrine Wnt signaling

The observed phenotypic differences between *Enl*-T1/*Wt1*^GFPCre^ and *Enl*-T1/*Six2*^GFPCre^ mouse models (**Figure 1** and **Figure S2**) could suggest a role for *Enl*-T1 stroma in contributing to developmental defects in the nephron. This prompted us to investigate whether *Enl*-T1 modulates the expression of genes involved in stroma-nephron interactions. We compiled a list of 28 genes previously shown or predicted to play a role in stroma-nephron interactions (Supplementary Table 13). Among them, *Pbx1*, *Meis1*, *Pdgfra*, and *Itga9* were down-regulated, while *Hoxa10*, *Hoxc10*, *Wnt5a,* and *Tcf21* were up-regulated in *Enl*-T1 *Foxd1*^+^ SP cells compared to *Enl*-WT (**Figure 6h, i**). Notably, *Pbx1* and *Meis1* are two major co-factors of *Hox*^74^. Genetic loss of *Pbx1* in the developing kidney results in smaller kidneys with fewer nephrons and expanded mesenchymal condensates in the nephrogenic area, akin to the phenotypes observed in *Enl*-T1 kidneys. The downregulation of these co-factors, coupled with upregulation of distinct *Hox* in *Enl*-T1 stroma, may result in the dysregulation of normal transcriptional programs governed by *Hox* TFs.

Among the genes upregulated in *Enl*-T1 *Foxd1*^+^ SP cells, *Tcf21*^75^ and *Wnt5a*^76^ are involved in the Wnt signaling pathway. RNA ISH experiments confirmed the upregulation of *Wnt5a* in stroma cells at the peripheral of CM (**Figure S9g**), likely representing *Foxd1*^+^ SP cells^71^. Aberrant activation of the Wnt pathway has been linked to impaired nephrogenesis and the onset of Wilms tumor^20,77^. *Tcf21* can interact with β-catenin and enhance the activation of its target genes^75^. Meanwhile, *Wnt5a* could act as an upstream ligand that binds to cell surface receptors to initiate the Wnt signaling cascade^78^. *Wnt5a* is normally expressed in the MS in the kidney (**Figure S9h**)^79^. Although Wnt5a typically transmits non-canonical Wnt pathway^80,81^, ectopic expression of *Wnt5a* can lead to abnormal activation of β-catenin/TCF signaling in calvarial mesenchyme in a transgenic mouse model^82^. Thus, the upregulation of *Wnt5a* and *Tcf21* in *Enl*-T1 *Foxd1*^+^ SP cells could potentially result in hyperactivation of the Wnt pathway in *Foxd1*^+^ SP cells. To test this hypothesis, we examined the expression of previously defined lineage-specific β-catenin-induced gene signatures^79^. Interestingly, we found that the expression of a nephron-specific, but not stroma-specific, β-catenin signature was elevated in *Enl*-T1 *Foxd1*^+^ SP cells compared to *Enl*-WT (**Figure S9i, j**). These results suggest that *Enl*-T1 *Foxd1*^+^ SP cells aberrantly express a β-catenin signature typically associated with the nephron lineage.

Previous studies have demonstrated that proper activation of β-catenin in NPCs is crucial for initiating mesenchymal-to-epithelial transition (MET) during nephrogenesis^77,83^, while its hyperactivation in *Six2^+^*NPCs or *Foxd1*^+^ SP cells can lead to an accumulation of undifferentiated nephron structures and a failure of MET^79,83^. Interestingly, we observed a marked increase in the nephron-specific β-catenin target gene signature in NP1, NP2, and T1-abnormal cells in *Enl*-T1 kidneys compared to their *Enl*-WT counterparts (**Figure 6j, k**). This elevation of β-catenin signature in *Enl*-T1 *Six2^+^* NPCs may be linked to an intrinsic increase in *Wnt4* (**Figure 4h**) and *Tcf21* (**Figure 5i**) in these cells. On the other hand, given the role of Wnt5a as a paracrine signal that activates the Wnt pathway in adjacent cells and the reported potential for stroma signals to amplify Wnt/β-catenin activity in NPCs^78,84^, the aberrant upregulation of *Wnt5a* in *Enl*-T1 *Foxd1*^+^ SP cells might also influence β-catenin activity in neighboring NPCs. Therefore, the nephrogenesis defects in *Enl*-T1 kidneys might be partially due to abnormal activation of the β-catenin/Wnt pathway in either or both the nephron and stroma (**Figure 6m**).

Next, we investigated the mechanism underlying ectopic upregulation of *Wnt5a* in *Enl*-T1 *Foxd1*^+^ SP cells. Visualization of ATAC intensity revealed that *Enl*-T1 *Foxd1*^+^ SP cells gained six DARs associated with the *Wnt5a* gene locus, with two in the promoter/genebody and four in distant regulatory elements (**Figure 6l**). There was a significant enrichment of *Hox9/11/12* motif sequences in all four distal DARs (**Figure S9k**), indicating a potential role of *Hox* TFs in the activation of *Wnt5a* through distal enhancers (**Figure 6m**).

### Blocking the acyl-binding activity of mutant ENL compromises its function on chromatin

As a chromatin reader, ENL binds to acylated histones to regulate transcriptional processes^29,30^. Although inhibitors designed to disrupt the acyl-binding activity of WT ENL proteins have been developed^85–91^, their effectiveness against ENL mutants has not been explored. Our previous work in HEK293 cells showed that ENL mutations found in Wilms tumor and AML (T1-T8) lead to enhanced self-association of ENL and the aberrant formation of condensates at select target genes, notably the *HOX* genes^33,34^. This leads to increased recruitment of ENL and its associated elongation factors, which in turn promotes the transcriptional elongation by RNA Polymerase II at these genes^34^. Disrupting condensate formation significantly impairs these mutants’ ability to activate target genes^34^. Furthermore, we found that disrupting the acyl-binding activity of these ENL mutants (T1, T1, T3) by introducing a point mutation (Y78A) in the YEATS domain results in a loss of their gene activation function^35,92^. These data imply that blocking the reader function of ENL mutants could be a viable strategy to inhibit their function.

We have developed a potent ENL inhibitor, TDI-11055 (**Figure 7a**), which effectively displaces WT ENL from chromatin by competitively binding to the acyl-binding pocket and successfully blocks its oncogenic function in AML^31^. Since cancer-associated mutations do not alter ENL’s acyl-binding pocket^35,92^, we hypothesized that TDI-11055 could also act on these mutants. In support, isothermal titration calorimetry assays confirmed the direct binding of TDI-11055 to purified YEATS domains harboring three different Wilms-tumor associated ENL mutations (T1, T2, T3)^35,92^, albeit with a slightly lower affinity than that observed with WT (**Figure 7b**). To assess the target engagement of TDI-11055 in a cellular context, we performed cellular thermal shift assays^93^ in HEK293 cells. TDI-11055 bound to and stabilized exogenously expressed Flag-tagged ENL-T1 and T2 proteins, but not to ENL-T1(Y78A) or ENL-T2 (Y78A) proteins that are deficient in binding acylated histones (**Figure 7c** and **Figure S10a**), supporting a direct interaction of this compound with ENL-T1/T2’s acyl-binding region.

**Figure 7.**
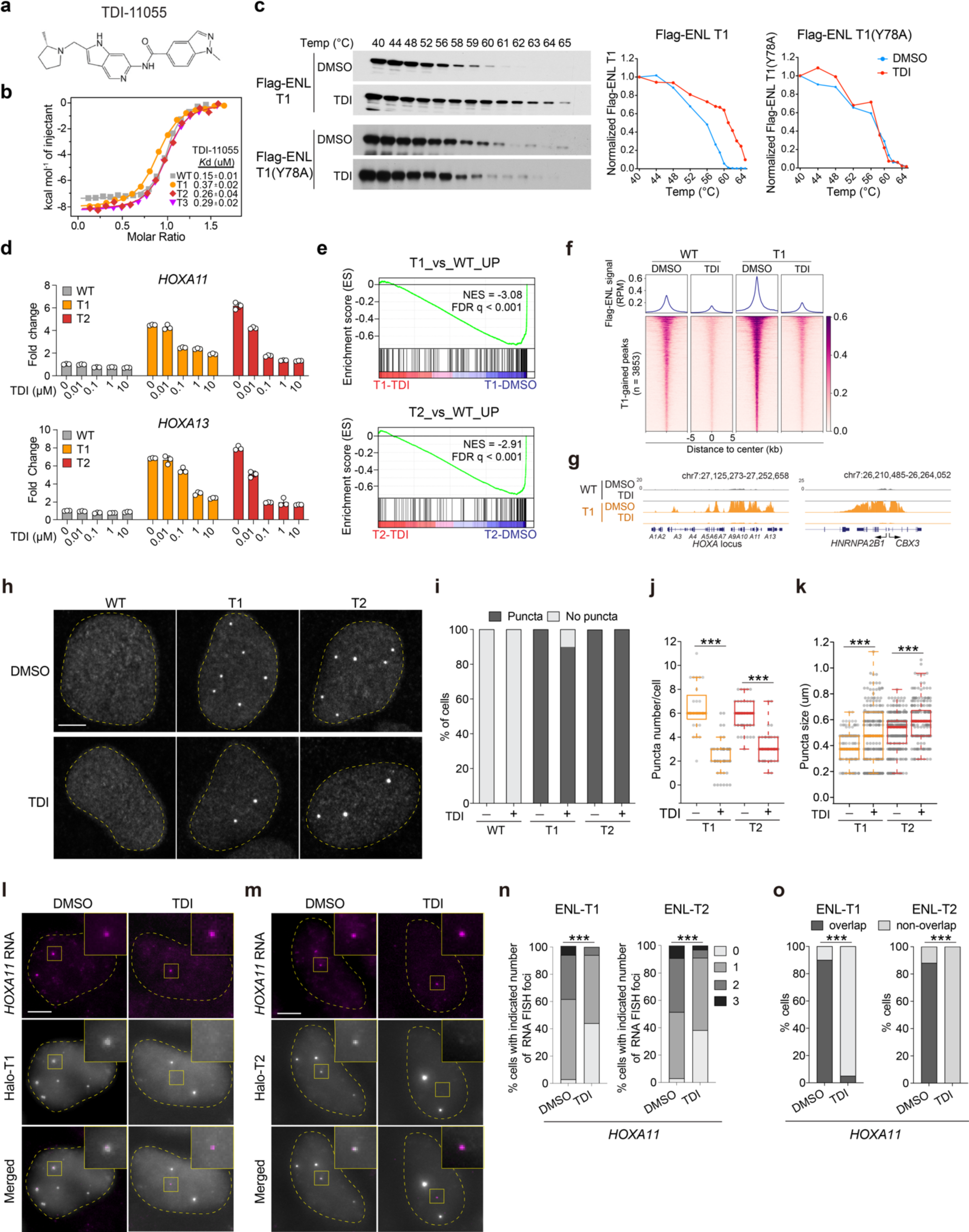
Blocking the acyl-binding activity of mutant ENL via a small-molecule inhibitor compromises its function on chromatin. **a**, Structure of the inhibitor, TDI-11055. **b**, ITC assay showing TDI-11055 directly binds to ENL-WT and ENL mutants (T1-T3) YEATS domains. **c**, Immunoblots and quantification showing the levels of Flag-ENL T1 and Flag-ENL T1(Y78A) after heat treatment in HEK293 cells at increasing temperatures. Temp, temperature. **d**, mRNA expression (normalized to *GAPDH*) of *HOXA* genes in HEK293 cells expressing endogenous levels of Flag-ENL transgenes with indicated treatment. DMSO was used as the vehicle. **e**, GSEA using custom gene set of upregulated genes in T1 or T2 vs. WT performed in cells treated with DMSO or TDI-11055. See Supplementary Table 14. **f**, NGS plot and heatmap of Flag-ENL ChIP-seq signals at up-bound peaks extracted from ENL-T1 DMSO vs. ENL-WT DMSO. See Supplementary Table 17. **g**, The genome browser view of Flag-ENL signals at select ENL-T1 target genes under DMSO or TDI-11055 treatment in HEK293 cells. **h**-**k**, IF staining of Flag-ENL (**h**) and quantification (**i**-**k**) in HEK293 cells under DMSO or TDI-11055 treatment. **i**, Percentage of nuclei with and without Flag-ENL condensates. **j**, the number of condensates in each nucleus. **k**, size of condensates. Center lines indicate median and box limits are set to the 25th and 75th percentiles. Quantifications performed on 29-52 cells. Two-tailed unpaired Student’s *t* test. \*\*\**P* < 0.001. **l**-**o**, Representative images of RNA-FISH (**l**, **m**) and quantification (**n**, **o**) showing the percentage of cells with indicated number of *HOXA11* nascent RNA FISH foci and the percentage of cells containing *HOXA11* nascent RNA FISH foci overlapped with Flag-ENL condensates (**o**). Chi-square test; \*\*\**P* < 0.001. **h**, **l**, **m**, Scale bar, 10 µm.

Next, we evaluated the effect of TDI-11055 on the transcriptional activity of ENL mutants in HEK293 cells. Treatment with TDI-11055 for 24 hours inhibited mutant ENL-induced increase in the expression of target genes *HOXA11* and *HOXA13* in a dosage-dependent manner (**Figure 7d**). To assess the global impact of TDI-11055 on mutant ENL-induced transcription, we performed RNA-seq on HEK293 cells treated with either DMSO or TDI-11055 for 24 hours. Transcriptional changes induced by TDI-11055 were more pronounced in ENL mutant cells compared to WT counterparts (**Figure S10b**). Gene set enrichment analysis (GSEA) revealed that genes upregulated by ENL mutants were strongly suppressed by TDI-11055 (**Figure 7e**). To investigate whether TDI-11055 inhibits mutant ENL-induced transcriptional changes by displacing it from chromatin, we performed chromatin immunoprecipitation followed by high-throughput DNA sequencing (ChIP–seq) for WT and T1 Flag-tagged-ENL in HEK293 cells treated with DMSO or TDI for 24 hours. ENL-T1 exhibited increased chromatin occupancy at a subset of target genes when compared with ENL-WT (**Figure 7f**), consistent with previous studies^35,92^. TDI-11055 treatment substantially decreased this enhanced chromatin binding by ENL-T1 at target genes (**Figure 7f**), such as *HOXA* and *CBX3* (**Figure 7g**). Together, these results demonstrate the efficacy of TDI-11055 in blocking mutant ENL’ chromatin binding and transcriptional function in cells.

We previously revealed that ENL mutants form submicron-sized condensates at specific genomic targets in HEK293 cells, and these condensates are functionally required for hyperactivation of these targets, including *HOXA11/13*^92^. Interestingly, we found that TDI-11055 treatment did not abolish the formation of mutant ENL condensates (**Figure 7h, i**), but rather decreased their number (**Figure 7j**) and slightly increased their size (**Figure 7k**). These effects phenocopy the acyl-binding defective Y78A mutation (**Figure S10c-l**). To understand how TDI-11055 impacts condensate localization to and expression of target genes, we performed IF staining for ENL mutants with concurrent nascent RNA FISH for *HOXA11* in HEK293 cells expressing Halo-tagged ENL-T1 (**Figure 7l**) or T2 (**Figure 7m**) proteins. TDI-11055 treatment reduced the number (**Figure 7n**) and intensity (**Figure S10m**) of *HOXA11* RNA FISH foci (**Figure 7n**), as well as the percentage of cells with one or more *HOXA11* FISH foci overlapping with a mutant ENL condensate (**Figure 7o**). Similar changes were not observed for a negative control gene, *GAPDH* (**Figure S10n-p**). These results suggest that TDI-11055 treatment could dislodge mutant ENL condensates from genomic targets, thereby abolishing gene activation induced by these condensates. Collectively, our data indicate that targeting the acyl-binding activity of ENL mutants via small molecules can abolish their function on chromatin and gene regulation in a cellular context.

### Transient *in vivo* treatment with TDI-11055 rescues mutant ENL-induced developmental defects

Building on the potent effect of TDI-11055 in cell lines, we next examined its potential to rescue mutant ENL-induced defects in kidney development. To this end, we treated pregnant mice with vehicle or TDI-11055 (100 mg/kg, once daily) via oral injection from E10.5, when mouse nephrogenesis begins^13^, to E14.5, followed by collection and histological characterization of kidneys on E15.5 (**Figure 8a**). Of note, such a short-term treatment with TDI-11055 did not alter the gross morphology (**Figure 8b**) nor cause noticeable changes in the histology of various nephron structures in *Enl*-WT kidneys (**Figure 8c-g**). Remarkably, TDI-11055 treatment restored the size of *Enl*-T1 kidneys to that of *Enl*-WT kidneys (**Figure 8b**). Histologically, abnormal structures observed in *Enl*-T1 kidneys, such as CM-UB structures and blastema-like structures, were markedly reduced upon TDI-11055 treatment (**Figure 8c**). Furthermore, TDI-11055 treatment partially rescued the decrease in differentiated nephron structures, including the Comma/S-shape bodies, glomeruli, and proximal and distal tubules (**Figure 8c-g**).

**Figure 8.**
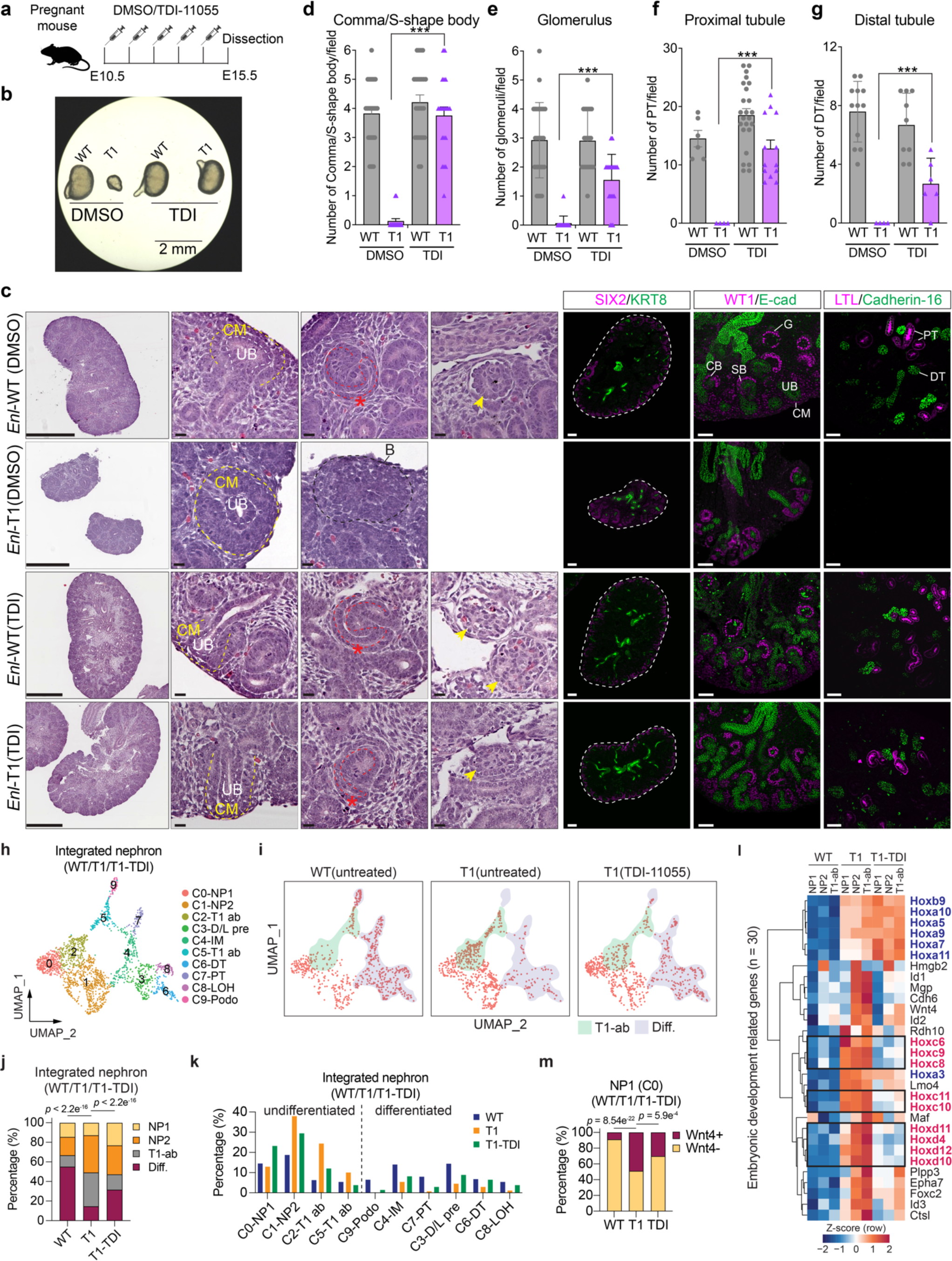
Transient *in vivo* treatment with TDI-11055 partially rescues mutant ENL-induced developmental and transcriptional defects in the developing kidney. **a**, Schematic to show the experimental dosing strategy. **b**, E15.5 kidneys. Scale bar, 2 mm. **c**, Left, Histology of E15.5 kidneys as described in **b**. The red star indicates S-shape body (SB) and the yellow arrows indicate glomerulus (G) structures. Scale bar in the first column images, 500 µm; scale bar in the zoom-in images, 20 µm. Right, Immunostaining for indicated proteins on E15.5 kidney sections. E-cad, E-cadherin. Scale bar in the first column images, 100 µm; scale bar in the zoom-in images, 50 µm. **d-g**, The number of nephron structures per field. **d**, Comma/S-shape body (Sample number of *Enl*-WT DMSO, *Enl*-T1 DMSO, *Enl*-WT TDI, and *Enl*-T1 TDI = 9, 10, 7, 6 kidneys); **e**, glomerulus (Sample number of *Enl*-WT DMSO, *Enl*-T1 DMSO, *Enl*-WT TDI, and *Enl*-T1 TDI = 7, 9, 7, 6 kidneys); **f**, proximal tubule (Sample number of *Enl*-WT DMSO, *Enl*-T1 DMSO, *Enl*-WT TDI, and *Enl*-T1 TDI = 4, 4, 7, 5 kidneys); **g**, distal tubule (Sample number of *Enl*-WT DMSO, *Enl*-T1 DMSO, *Enl*-WT TDI, and *Enl*-T1 TDI = 4, 4, 3, 3 kidneys). One dot indicates the number of one indicated structure per field. Date represent mean ± s.d.; two-tailed unpaired Student’s *t*-test, \*\*\**P* < 0.001. **h, i**, UMAP embedding of integrated scRNA-seq cells from *Enl*-WT, T1, and T1-TDI nephrons. **h**, Cells are colored and labeled by annotated cell types. **i**, T1-ab clusters (C2/5) are highlighted in green; Differentiated structures (C3/4/6/7/8) are highlighted in purple. Diff., differentiated. **j**, The percentage of main nephron cell types within *Enl*-WT, T1, and T1-TDI nephrons. Chi-Square test *p*-values are shown. **k**, The percentage of all nephron clusters within *Enl*-WT, T1, and T1-TDI nephrons. **l,** The expression of embryonic development-related T1-UP DEGs identified in Figure 2k in NP1, NP2, and T1-ab cells from *Enl*-WT, T1, and T1-TDI nephrons. Gene expression is normalized by Z-score. *Hox* genes rescued by TDI-11055 treatment are highlighted in red text and those not rescued are highlighted in blue. **m**, The percentage of NP1 cells with or without *Wnt4* expression in *Enl*-WT, T1, or T1-TDI nephrons. Fisher’s exact test *p*-values are shown.

To elucidate the cellular and molecular changes induced by TDI-11055 at single-cell resolution, we performed scRNA-seq on E15.5 *Enl*-T1 kidneys following treatment with TDI-11055 (**Figure S11a, b**). By integrating scRNA-seq datasets from untreated *Enl*-WT and *Enl*-T1 kidneys, along with *Enl*-T1 kidneys treated with TDI-11055, we classified all cells into four distinct lineages and assessed their distribution. Our analysis revealed that TDI-11055 treatment largely reverted T1-induced alterations in the proportions of nephron and stroma cells (**Figure S11c**). Next, we identified distinct clusters in the nephron compartment representing 10 different cell types (**Figure 8h** and **Figure S11d**). Compared to untreated *Enl*-T1 kidneys, TDI-11055-treated *Enl*-T1 kidneys showed a higher representation of various differentiated structures and a lower representation of T1-abnormal populations (C2 and C5) (**Figure 8i-k**). Moreover, TDI-11055 treatment suppressed a significant portion of genes upregulated by mutant ENL in nephron progenitor clusters (NP1, NP2, T1-ab). Notably, this included several *Hox* genes, particularly *Hoxc* and *Hoxd* genes, as well as *Wnt4* (**Figure 8l**). TDI-11055 treatment also partially reverted the increase in the percentage of *Wnt4^+^* cells in *Enl*-T1 NP1 cells (**Figure 8m**). In contrast, TDI-11055 treatment did not affect the expression levels of T1-upregulated genes associated with mitochondrial and metabolic pathways (**Figure S11e**). For T1-downregulated DEGs in the nephron, TDI-11055 treatment only partially restored the expression levels of a small subset of genes (**Figure S11f**). These included *Six2* and *Cited1* in *Enl*-T1 NP1 (**Figure S11g**), suggesting that the self-renewal of *Enl*-T1 NPCs is partially recovered. Importantly, TDI-11055 treatment reduced the transcriptomic similarity of *Enl*-T1 nephron progenitor subsets to *ENL*-mutant Wilms tumor (**Figure S11h**). Furthermore, TDI-11055 treatment reduced *Wnt5a* expression levels in *Enl*-T1 *Foxd1*^+^ SP cells (**Figure S11i**). The β-catenin activation signature was downregulated by TDI-11055 treatment in both nephron (NP1, NP2, T1-ab) and stroma (*Foxd1*^+^ SP) (**Figure S11j**). Altogether, these results demonstrate that transient inhibition of the acyl-binding activity can partially rescue mutant ENL-induced transcriptomic and developmental alterations *in vivo*.

## Discussion

Despite the established link between disrupted development and cancer, our understanding of how specific cancer mutations impact gene regulatory landscapes to impair developmental programs *in vivo* remains rather limited. Our study provides new insights into this fundamental question by focusing on ENL, the most frequently mutated epigenetic regulator in Wilms tumor^20^. Through the integration of genetic mouse modeling, histological characterizations, and single-cell analyses, we have uncovered the role of the *ENL*-T1 mutation in perturbing kidney development and elucidated the underlying mechanisms at single-cell resolution (**Figure 9**). We have also demonstrated that transient inhibition of the chromatin reader activity of mutant ENL can effectively reverse these alterations, thus presenting a proof-of-concept for the potential use of epigenetics-targeted agents in correcting developmental defects.

**Figure 9.**
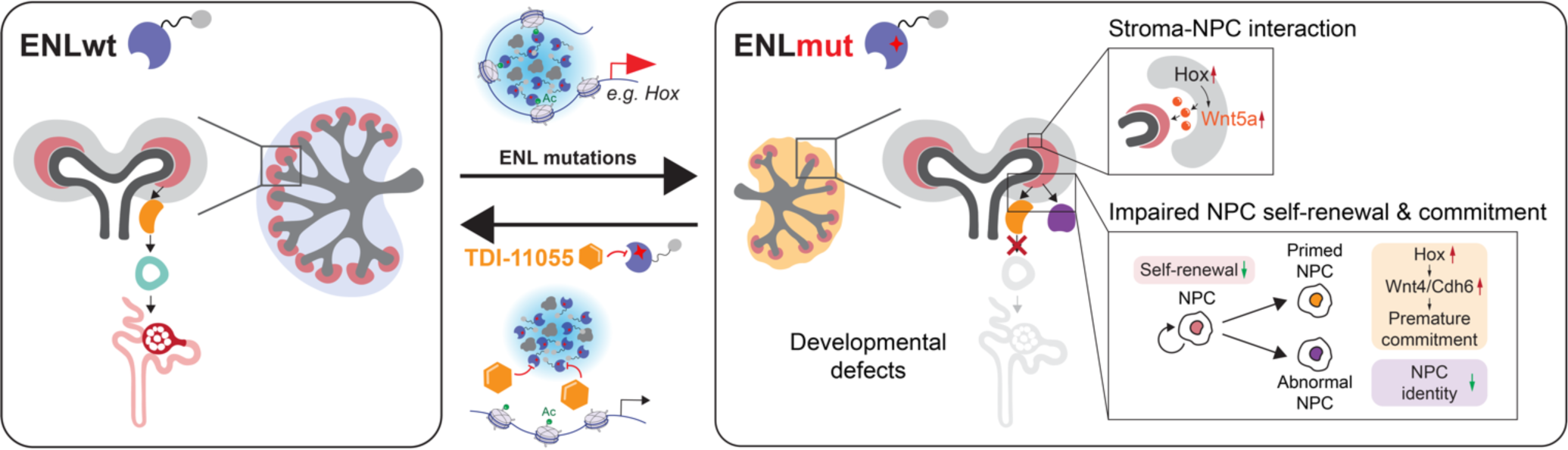
Overall summary of the study. Schematic showing normal nephrogenesis (left box) and that ENL mutation impairs kidney development trajectory by rewiring gene regulatory landscape (right box). The mutant ENL (ENLmut) disrupts kidney development by driving nephron progenitors (NPC) into a committed state while concurrently impeding their further differentiation. This dysregulation involves the misregulation of critical transcription factor regulons, particularly the *HOX* clusters. ENLmut forms transcriptional condensates at *HOX* clusters and hyper-activates *HOX* genes, which in turn, leads to increased expression of priming factors such as *Wnt4* and *Cdh6* in NPCs. Additionally, ENLmut induces the emergence of abnormal NPCs that lose the chromatin identity typically associated with kidney development. Furthermore, ENLmut might disrupt stroma-nephron interactions through hyperactivation of paracrine Wnt5a signaling. These multifaceted effects resulting from the mutation lead to severe developmental defects in the kidney and early postnatal mortality in mice. Inhibition of the acetylation binding activity of ENLmut with a small molecule (TDI-11055) displaces ENLmut condensates from target genes and abolishes its gene activation function. This intervention effectively restores developmental defects in mice.

Our integrated single-cell transcriptomics and chromatin accessibility profiling produced a cell atlas of mouse embryonic kidneys, elucidated key regulatory mechanisms for each cell type, and revealed dynamic changes in open chromatin accessibility during cell fate transitions. This work, in conjunction with previous studies that applied similar technologies to late-stage developing and adult kidneys, offers a comprehensive map for understanding normal kidney function and disease development. Building on this information, we demonstrated that a Wilms tumor-associated ENL mutation (T1) altered the cellular state and composition of the nephron. Specifically, *Enl*-T1 causes an accumulation of nephron progenitor populations and a reduction of various differentiated cell types, indicating a block in renal differentiation. Intriguingly, *Enl*-T1 nephron progenitors (NP1) exhibit a more committed state, as evident by their chromatin accessibility landscapes and marker gene expression (e.g., downregulation of *Six2* and *Cited1* and upregulation of *Wnt4* and *Chd6*). While it has been proposed that nephron progenitors in Wilms tumor display elevated self-renewal ability^94^, our findings highlight unexpected effects of cancer mutations on the balance of NPC self-renewal and commitment during nephrogenesis. Additionally, we observed the emergence of abnormal, poorly differentiated cells (T1-ab) in *Enl*-T1 nephrons with distinct transcriptomic and chromatin profiles compared to self-renewing (NP1) and committed (NP2) progenitors. These cells maintain a progenitor-like state but lose the potential for kidney specification, possibly due to a decrease in *Six2* expression and/or activity. Furthermore, compared to committed nephron progenitors (NP2), these cells exhibit a severe defect in proliferation, a property critical for NPC commitment and differentiation^13,14,95^. Future studies are warranted to investigate whether other Wilms tumor-associated mutations also induce such a de novo cell state in the developing kidney and whether these cells contribute to Wilms pathogenesis.

The developmental defects observed in *Enl*-mutant kidneys can be attributed to specific alterations in the transcriptome and chromatin accessibility. Notably, mutant ENL upregulates multiple *Hox* clusters across various cell types, including nephron progenitors (NP1, NP2, and T1-ab) and *Foxd1*^+^ stroma progenitors. This, along with our previous studies in HEK293 cells showing direct binding of ENL mutants to *HOX* genes^33,34^, establish *HOX* genes as direct targets of ENL mutants. Moreover, *Hox* genes are identified as top TF candidates responsible for the changes in chromatin accessibility induced by mutant ENL in nephron and stroma progenitors. These results suggest a critical role for *Hox* genes and their downstream targets, such as *Wnt4* and *Cdh6,* in mediating kidney phenotypes in *Enl-T1* mice. Future studies should explore whether decreasing *Hox* gene expression in *Enl*-T1 kidneys can mitigate the nephrogenesis defects.

Among the *Hox* genes upregulated in *Enl*-T1 kidneys, the genetic loss of *Hox9/10/11* has been reported to result in lineage infidelity in the kidney^96^, highlighting the role of these genes for proper lineage specification and maintenance. Our study suggests that persistent hyperactivation of certain *Hox* genes could promote immature commitment of NPCs while simultaneously restricting them from further differentiation. Importantly, *ENL*-mutant Wilms tumors express higher levels of certain *HOX* genes compared with *ENL*-WT tumors^32^. These findings suggest that maintaining proper expression levels of *HOX* genes is essential for the precise execution of kidney differentiation. Dysregulation of *HOX* genes caused by mutations in *ENL* and potentially other Wilms tumor-associated genes may contribute to Wilms tumor pathogenesis, an area that warrants further investigation.

Our study suggests a potential role for mutant ENL in perturbing the normal stroma-nephron interaction critical for nephrogenesis^17^. Specifically, *Enl*-T1 expression leads to substantial alterations in the open chromatin landscape of *Foxd1*^+^ stroma progenitors, which locate adjacent to the CM and are known to play a vital role in stroma-nephron interactions. Several genes involved in stroma-nephron interactions^73,75,76^, such as *Hoxa10*, *Hoxc10*, *Wnt5a*, and *Tcf21*, are upregulated in *Enl*-T1 *Foxd1*^+^ SP cells. *Wnt5a* and *Tcf21* are linked to the Wnt signaling pathway. Proper activation of the Wnt pathway in both stroma and nephron is required for balanced NPC self-renewal and differentiation^51^. It is tempting to speculate that *Enl*-T1 *Foxd1*^+^ SP might contribute to the aberrant activation of the Wnt pathway observed their adjacent NPCs by secreting de novo-gained *Wnt5a*, thereby influencing renal differentiation. Future studies are needed to thoroughly test this hypothesis. Interestingly, several enhancers associated with the *Wnt5a* locus display increased chromatin accessibility in *Enl*-T1 *Foxd1*^+^ SP cells and contain *Hoxa* binding sites, offering a potential mechanism for enhanced *Wnt5a* expression in these cells. Notably, *CTNNB1* is one of the most frequently mutated genes in Wilms tumors, and up to 50% of Wilms tumors exhibit nuclear accumulation of β-catenin indicative of constitutive activation of the β-catenin pathway^79^. Therefore, ENL mutations may represent a previously unknown mechanism to activate this pathway in the developing kidney. Unbiased grouping of human Wilms tumors based on their transcriptomes results in multiple clusters, with *ENL*, *CTNNB1*, and *WT1* mutations residing in similar clusters^20,32^, supporting a molecular link between these proteins in kidney biology and diseases. Thus, insights gained from studying *ENL* mutations could have implications for a significant proportion of Wilms tumors.

Wilms tumors are highly heterogenous in histology and contain cells from diverse lineages, including the nephron, stroma, and muscle^5,10–12,37^. Thus, it remains a challenge to pinpoint cell types and/or developmental states that are susceptible to transformation. By utilizing the *Wt1*-cre strain to induce *Enl*-T1 expression in early progenitors that give rise to both nephron and stroma lineages, we have revealed the role of this mutant in remodeling transcriptional and chromatin accessibility landscapes in within these two compartments. Future studies are needed to determine whether mutant ENL can also affect other lineages within the kidney, such as the ureteric bud/ureteric epithelium (UB/UE) and endothelial cells. Nevertheless, our study may suggest that the nephron and stroma lineages or their early progenitors are potential cellular targets for Wilms tumor-associated mutations. Lineage-specific investigations^21–23,79^ of several Wilms tumor-associated mutations support this hypothesis. For instance, gain-of-function β-catenin mutations have been detected in both the blastema and stromal components of human Wilms tumor^97^. While earlier studies suggest that β-catenin activation in the nephron is causal to Wilms tumor formation^23,98^, recent work demonstrates that activation of β-catenin specifically in the stroma in mouse models non-autonomously prevents the NPC differentiation and results in histological and molecular features that resemble human Wilms tumors^79^. Furthermore, the overexpression of *Lin28*, a gene encoding for an RNA-binding protein and amplified in Wilms tumor, results in Wilms tumors formation in mice only when introduced into early renal progenitors that give rise to both nephron and stroma lineages^21^. Given the histological and molecular diversity observed in Wilms tumors, it is conceivable that they differ in their developmental roots and mechanisms. As exemplified in our study of ENL mutations, the integration of genetic mouse modeling and single-cell technologies holds the promise of elucidating these fundamental questions, thereby advancing our basic understanding of kidney development and pathogenesis at the cellular and molecular levels.

Recent genomics studies have revealed that a significant number of mutations identified in Wilms tumor impact genes associated with histone modification and transcriptional elongation. These genes include *ENL (MLLT1)*, *BCOR*, *BCORL1*, *HDAC4*, *EP300*, *CREBBP*, *BRD7*, and *MAP3K4*^20^. Moreover, germline mutations that predispose individuals to Wilms tumor are also found in genes that converge into similar pathways, such as *CDC73* and *CTR9*^20^. Notably, ENL has been shown to interact with several of these factors, including BCOR^99^, CDC73 and CTR9^100^, suggesting potential functional interconnectedness. Our work showcases the significant impact of mutated epigenetic regulators on kidney development, serving as a catalyst for future investigations into other epigenetic factors. A limitation of our study, however, is that the profound development defects and early lethality caused by mutant ENL expression in *Wt1*^+^ precursors preclude a direct assessment of its role in Wilms tumor formation, which typically manifests months after birth. Given that ENL mutations often occur in Wilms tumor without other well-known genetic alterations, except in occasional instances with *CTNNB1* mutations^32^, future studies should consider inducing ENL mutations, either alone or in combination with *CTNNB1* mutations, in a limited subset of kidney progenitors. This strategy would better recapitulate the sporadic nature of mutational events in human cancer and likely circumvent the detrimental developmental effects observed in the *Wt1*-Cre/*Enl*-T1 model, thus allowing for a more targeted exploration of the potential role of ENL mutations in tumorigenesis.

Given the amenability of epigenetic mechanisms for therapeutic intervention, research in this area holds promise for uncovering novel treatment avenues. We have demonstrated that a small molecule inhibitor specifically targeting the acyl-binding activity^31^ can rescue developmental and transcriptional defects induced by mutant ENL in embryonic kidneys. This serves as a compelling example of how targeting epigenetic mechanisms can mitigate developmental abnormalities caused by disease mutations. Our results also imply that inhibitors of the ENL YEATS domain may be promising therapeutic agents for treating cancers driven by these ENL mutations. However, given that these inhibitors also affect the wildtype ENL, a careful assessment of potential side effects is essential if this strategy is to be translated into clinical setting. Moreover, as exemplified by the study of EZH2 gain-of-function mutations in lymphoma^101^, genetic mutations in specific genes may imply hitherto undiscovered functions of wildtype proteins in the same disease context. Future studies are needed to explore the potential involvement of wildtype ENL in kidney development and Wilms tumors lacking ENL mutations, as well as to explore the broader applicability of ENL inhibitors.

In summary, we have provided a direct examination of the molecular and cellular consequences of an ENL mutation associated with Wilms tumor during kidney development. These studies identified key epigenetic and transcriptional aberrations that disrupt cellular differentiation programs *in vivo* and may contribute to disease. Our work also highlights the power of combining genetic mouse models and emerging single-cell multi-omics approaches in understanding how disease mutations perturb normal development and exploring potential therapeutic avenues.

## Data Availability

Raw data, processed data, and metadata from mouse single-cell datasets have been deposited in GEO with the accession number GSE243868 and GSE243870. The ChIP-seq and RNA-seq data have been deposited in the Gene Expression Omnibus database under accession numbers GSE243866 and GSE243867. All other raw data generated or analyzed during this study are included in this published article (and its Supplementary Information files). Codes used for data analysis are available upon request.

## Acknowledgments

The authors thank members of the Wan laboratory for scientific input throughout the study; Dr. Kai Tan for critical reading of the manuscript; Dr. Xianyang Deng for scientific input; Dr. Yuting Guan and Dr. Yuka Sakata for technical support; Krista A. Budinich, Sylvia Tang, and Chujie Gong for manuscript editing; the CDB Microscopy Core and ULAR facility at the University of Pennsylvania for facility support. The research was supported by funds from the University of Pennsylvania (L.W.), a NIH Director’s New Innovator Award (1DP2HG012443 to L.W.), a NIH Pathway to Independence Award (R00CA226399 to L.W.), and a Pew-Stewart Scholar Award (L.W.). L.W. is a scholar of The Leukemia & Lymphoma Society.

## Author contributions

L.W., L.S., and Q.L. conceived and designed the overall study. L.S., L.X., and Q.L. performed mouse works; L.S. performed cellular studies; Q.L. performed bioinformatic data analysis. Y.L. and H.L. performed ITC assay. H.W. and Q.Q. provided support for scRNA-seq and snATAC-seq studies; L.W., L.S., and Q.L. wrote the manuscript with critical input from K.S., H.W., K.S., and Q.Q.. L.W. supervised the overall study.

## Supplementary

**Figure S1.**
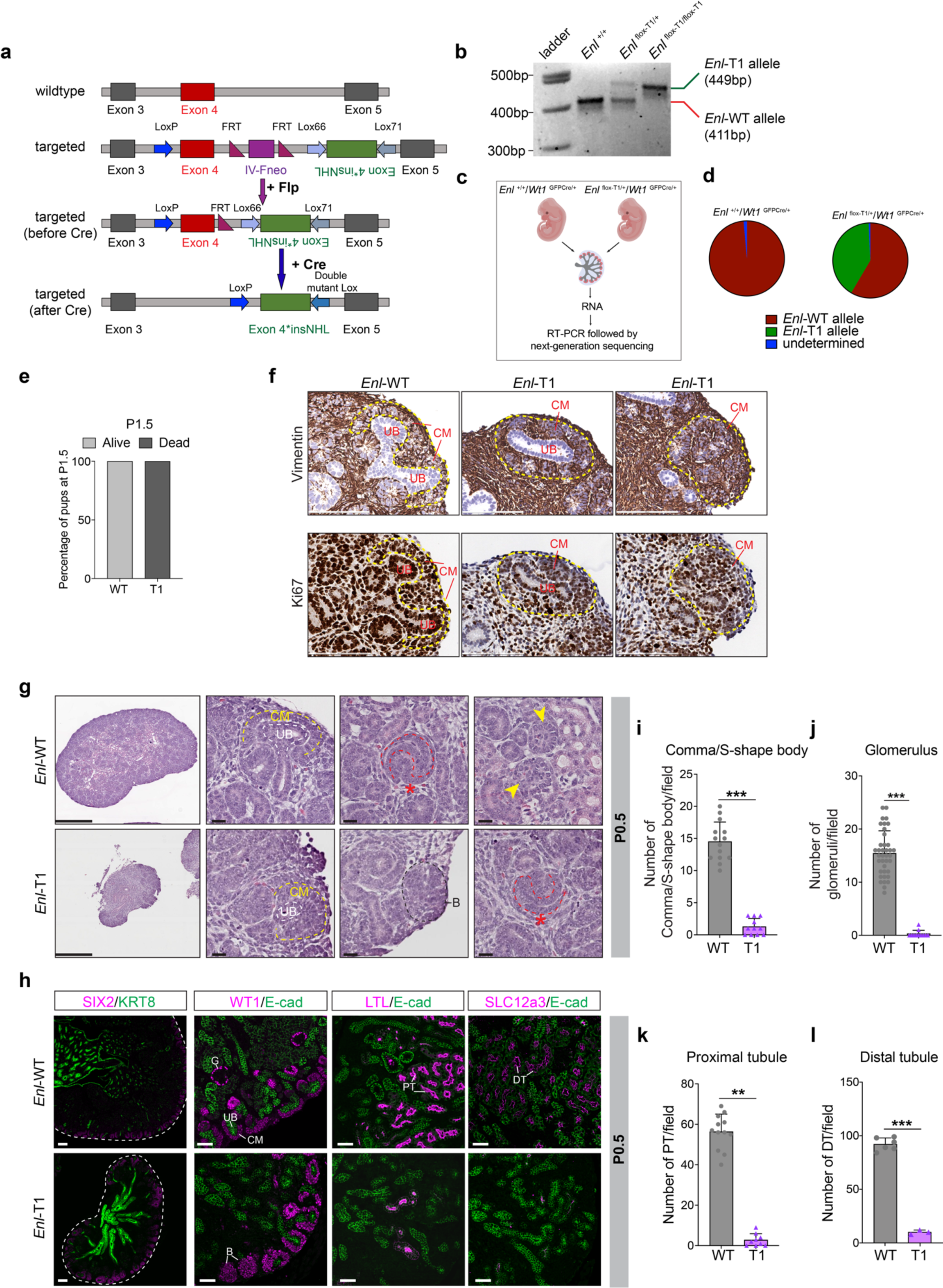
Heterozygous expression of mutant ENL disrupts embryonic kidney development and leads to postnatal mortality in *Wt1*^GFPCre/+^ mice. **a**, Schematic of the *Enl*-T1 knock-in mouse model design. **b**, Gel image of genotyping PCR for indicated mice. **c**, Schematic of confirming the expression of *Enl*-T1 in kidney upon Cre-recombinase. **d**, The percentage of *Enl*-WT and *Enl*-T1 mRNA detected in E15.5 kidney. **e**, Quantification of survival status of *Enl*-WT and *Enl*-T1 pups at P1.5. **f**, Representive images of Ki-67 and Vimentin staining in E15.5 *Enl*-WT and *Enl*-T1 kidneys. Scale bar = 100 µm. **g**, Hematoxylin and eosin–stained sections showing the histology of P0.5 kidneys from *Enl*-WT and *Enl*-T1 pups. CM, cap mesenchyme; UB, ureteric bud. The red star indicates S-shape body, the black dashed line outlines a region of blastema-like structure (B), and the yellow arrows indicate glomerulus structures. Scale bar in the first column images, 500 µm; scale bar in the zoom-in images, 20 µm. **h**, Immunostaining for SIX2, KRT8, WT1, E-cadherin (E-cad), LTL, and SLC12a3 on P0.5 kidney sections. Scale bar in the first column images, 100 µm; scale bar in the zoom-in images, 50 µm. CM, cap mesenchyme; UB, ureteric bud; B, blastema-like structure; G, glomerulus; PT, proximal tubule; DT, distal tubule. **i**-**l**, Number of nephron structures per field. **i**, Comma/S-shape body (*n* = 8 *Enl*-WT kidneys and 6 *Enl*-T1 kidneys); **j**, glomerulus (*n* = 11 *Enl*-WT kidneys and 6 *Enl*-T1 kidneys); **k**, proximal tubule (*n* = 5 *Enl*-WT kidneys and 6 *Enl*-T1 kidneys); **l**, distal tubule (*n* = 4 *Enl*-WT kidneys and 4 *Enl*-T1 kidneys). One dot indicates the number of one indicated structure per field. Date represent mean ± s.d.; two-tailed unpaired Student’s *t*-test, \*\*\**P* < 0.001.

**Figure S2.**
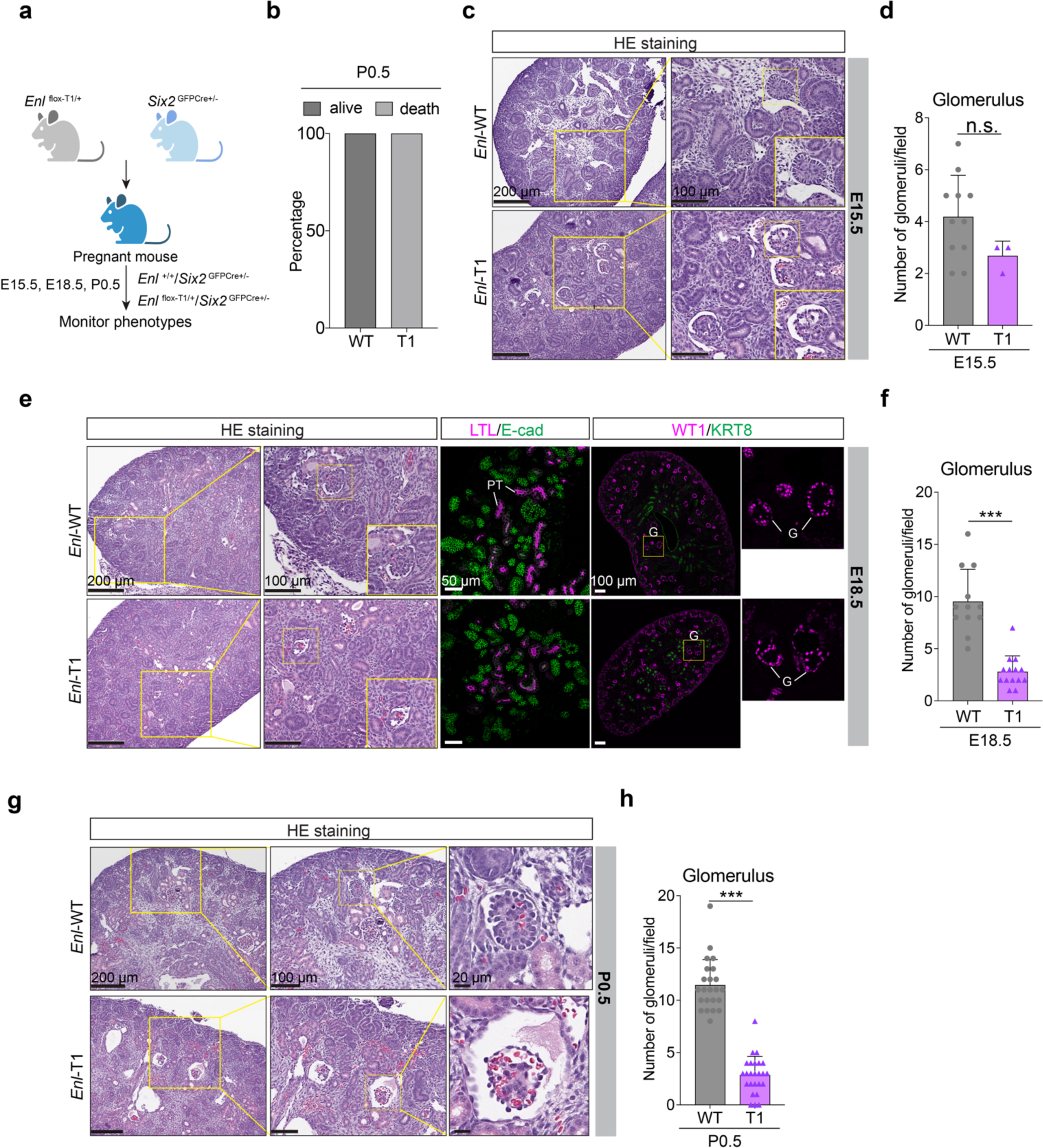
Heterozygous expression of mutant ENL disrupts embryonic kidney development and leads to postnatal mortality in *Six2*^GFPCre/+^ mice. **a**, Schematic of the breeding strategy. **b**, Quantification of survival status of *Enl*-WT and *Enl*-T1 pups at P0.5. **c**, **e**, **g**, Hematoxylin and eosin– stained sections showing the histology of E15.5 (**c**), E18.5 (**e**), and P0.5 (**g**) kidneys from *Enl*-WT and *Enl*-T1 pups. Scale bars are labeled in the images. **e**, Immunostaining for WT1, KRT8, E-cadherin (E-cad), and LTL on E18.5 kidney sections. Scale bars are labeled in the images. G, glomerulus; PT, proximal tubule. **d, f, h**, Number of glomeruli per field (E15.5, *n* = 6 *Enl*-WT kidneys and 2 *Enl*-T1 kidneys; E18.5, *n* = 4 *Enl*-WT kidneys and 6 *Enl*-T1 kidneys; P0.5, *n* = 8 *Enl*-WT kidneys and 9 *Enl*-T1 kidneys). One dot indicates the number of one indicated structure per field. Date represent mean ± s.d.; two-tailed unpaired Student’s *t*-test, \*\*\**P* < 0.001. n.s., no significance

**Figure S3.**
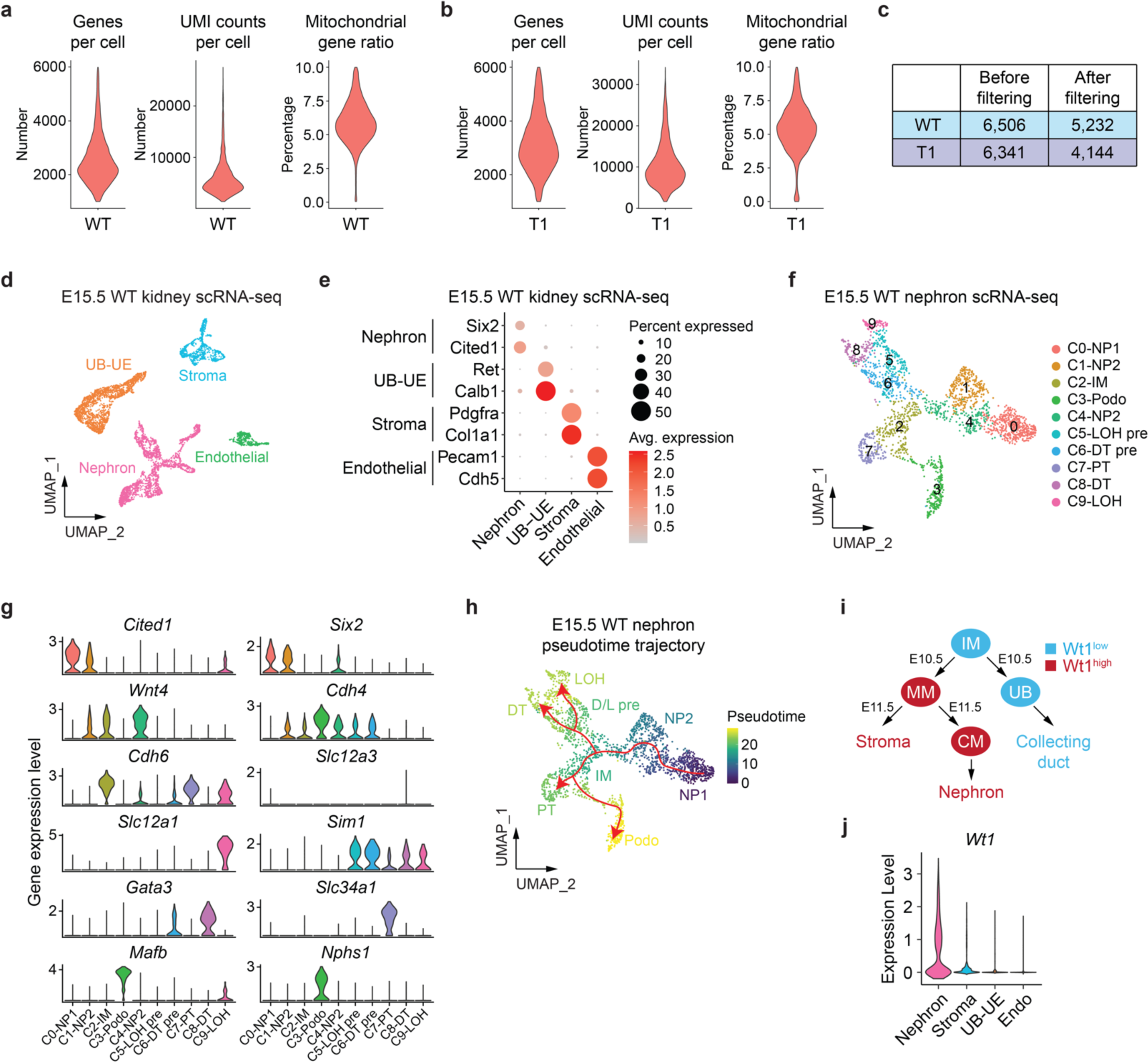
scRNA-seq identifies major cell types in *Enl*-WT embryonic kidneys. **a**, **b**, Violin plot showing the number of informative genes per single cell, unique molecular identifiers (UMIs) per single cell, and mitochondrial gene ratio in scRNA-seq dataset of *Enl*-WT (**a**) and T1 (**b**) kidneys after quality control filtering. **c**, Table showing the number of cells before and after filtering according to the criteria written in Methods. **d**, UMAP embedding of *Enl*-WT kidney scRNA-seq data, with cells colored to represent the four main embryonic kidney lineages. UB, ureteric bud; UE, ureteric epithelium. **e**, Dot plot showing the expression of selected marker genes for each main kidney lineage. Color scale represents the average (Avg.) expression level. Circle size represents the percentage of cells expressing the gene. **f**, UMAP embedding of integrated scRNA-seq cells from *Enl*-WT nephrons. Cells are colored and labeled by annotated cell types. NP, nephron progenitor; IM, intermediate stage; Podo, podocyte; LOH pre, loop-of-Henle precursor; DT pre, distal tubule precursor; PT, proximal tubule; DT, distal tubule; LOH, loop-of-Henle. **g**, Violin plot showing the gene expression of selected makers for each cell type in *Enl*-WT nephrons. **h**, UMAP embedding of *Enl*-WT nephron scRNA-seq differentiation trajectory. Cells are colored by pseudotime. Trajectory is depicted by red arrows. **i**, Schematic illustrating the dynamics of *Wt1* gene expression in different cell lineages during nephrogenesis. Lowly and highly *Wt1* expressing lineages were colored in blue or red, respectively. IM, intermediate mesoderm; MM, metanephric mesenchyme. **j**, Violin plot showing the gene expression of *Wt1* in indicated cell lineages from *Enl*-WT kidneys. Endo, endothelial cells.

**Figure S4.**
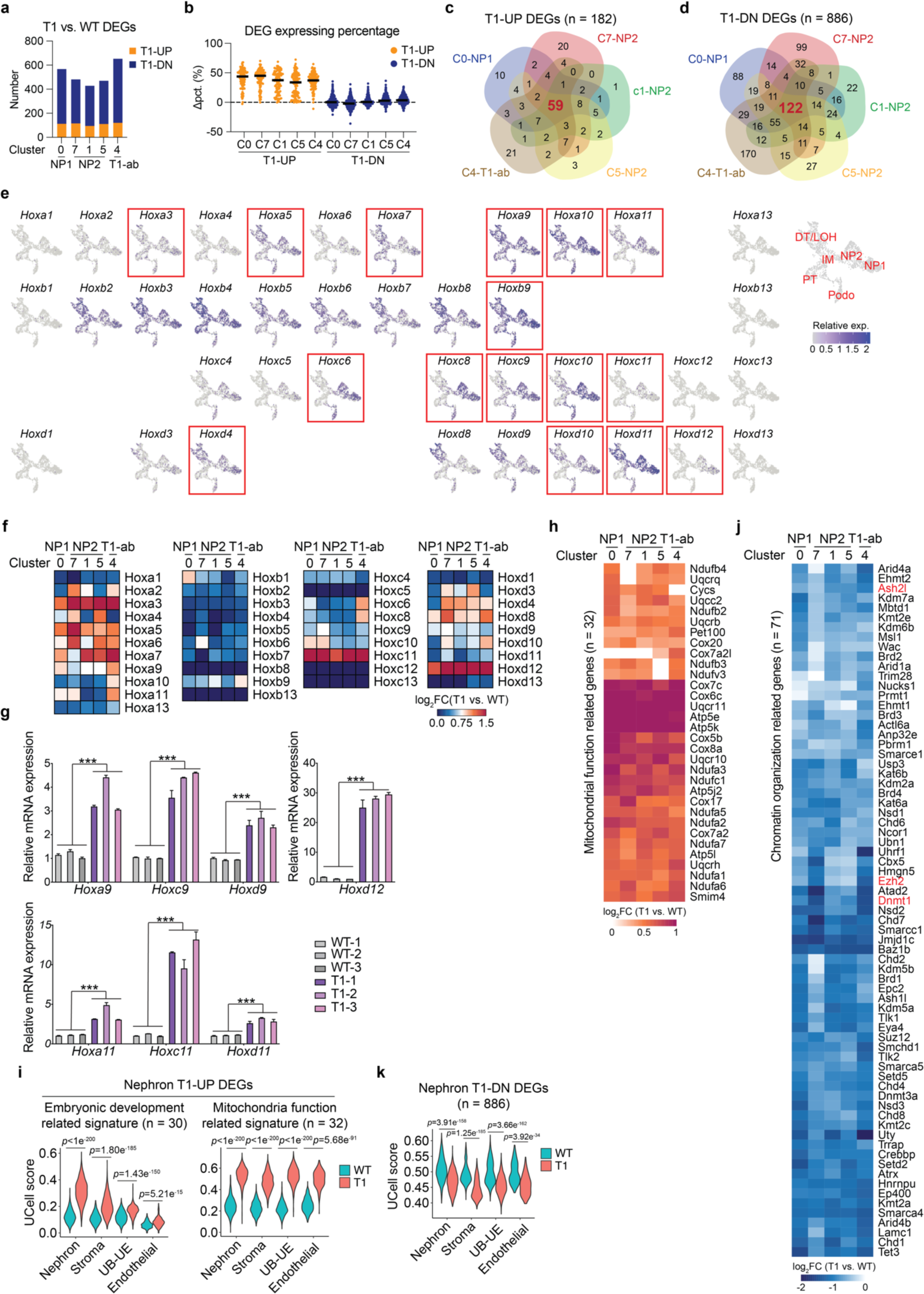
Transcriptional changes induced by mutant ENL in the developing kidney. **a**, Stacked bar plot showing the number of DEGs between *Enl*-WT and T1 (upregulated in T1, T1-UP; downregulated in T1, T1-DN) in indicated nephron cell types. **b**, Dot plot showing the differential expressing percentage (Δpct.) of DEGs between *Enl*-WT and T1 in indicated nephron cell types. Δpct. = pct. (*Enl*-T1) - pct. (*Enl*-WT). T1-UP and DN DEGs were colored in orange or blue, respectively. **c**, **d**, Venn diagram showing the overlap among cell type specific T1-UP (**c**) or T1-DN (**d**) DEGs. **e**, UMAP embeddings showing all 39 *Hox* gene expression patterns in the *Enl*-WT nephron. Cells are colored by the expression level. The *Hox* genes that appeared in the union T1-UP DEG list were highlighted with a red box. Nephron cell types are labeled on the right. **f**, Heatmap showing the gene expression fold change (*Enl*-T1 versus WT) of all 39 *Hox* genes in indicated cell types. Fold change is scaled by log2. **g**, mRNA expression levels of *Hox* genes (normalized to *GAPDH*) in E15.5 *Enl*-WT and *Enl*-T1 kidneys. \*\*\**P* < 0.001. **h**, Heatmap showing the fold change of mitochondrial and metabolism-related T1-UP DEGs identified in Figure 2i within the indicated nephron cell types. Fold change is scaled by log2. **i**, **k**, Violin plot showing the UCell score evaluated by embryonic development-related (**i**, left), mitochondrial and metabolism-related (**i**, right), and union T1-DN (**k**) signatures for the four main embryonic kidney lineages within *Enl*-WT and T1 kidneys. Wilcoxon rank-sum test *p*-values are shown. **j**, Heatmap showing the fold change of chromatin organization-related T1-DN DEGs identified in Figure 2m within the indicated nephron cell types. Fold change is scaled by log2. Epigenetic regulators Dnmt1, Ezh2, and Ash2l are highlighted in red.

**Figure S5.**
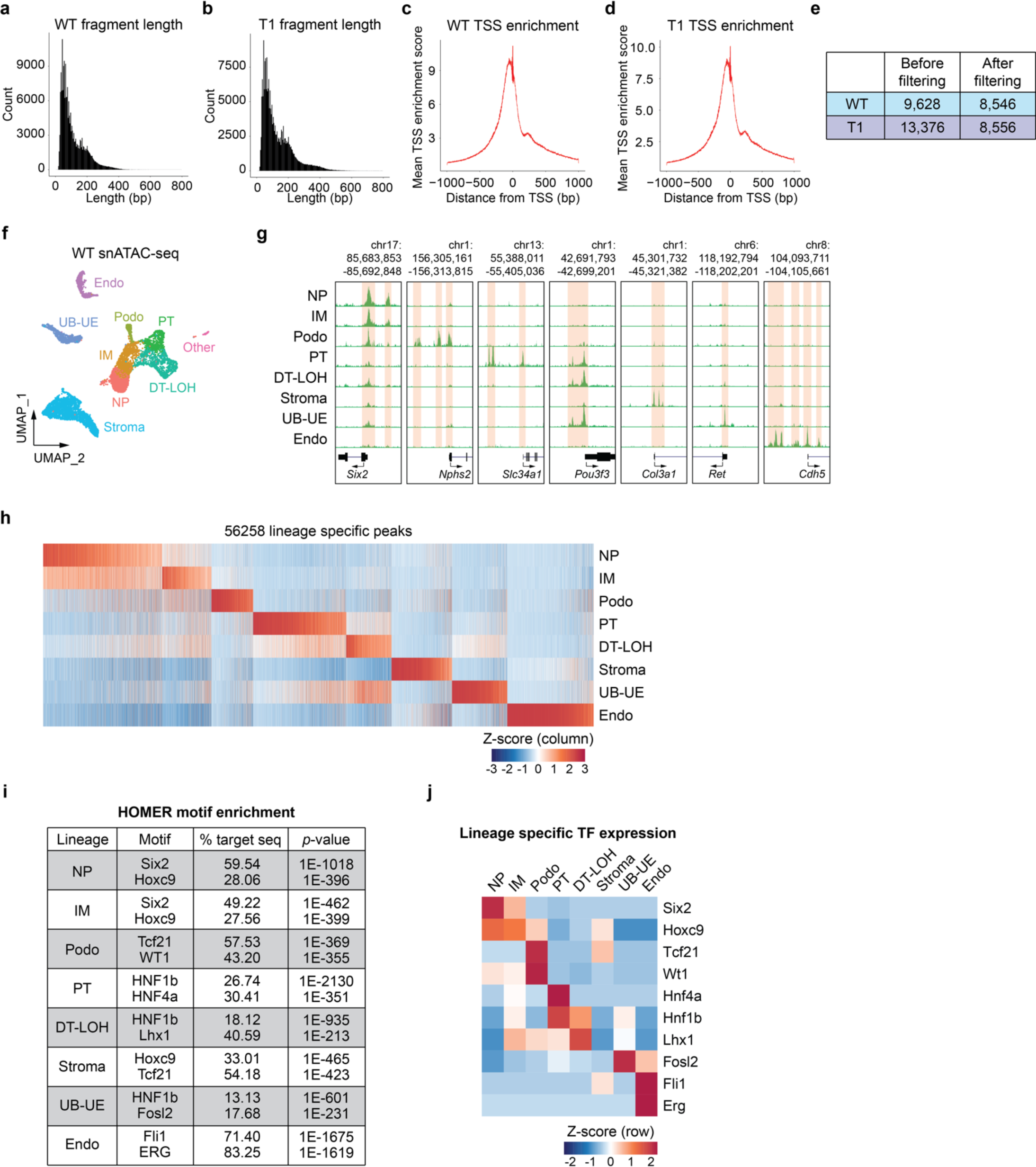
snATAC-seq reveals cell-type specific regulators in E15.5 *Enl*-WT kidney. **a**, **b**, Insert size distribution of *Enl*-WT (**a**) and T1 (**b**) snATAC-seq data showing periodic patterns. **c**, **d**, Transcription start site (TSS) signal enrichment of the *Enl*-WT (**c**) and T1 (**d**) snATAC-seq data. **e**, Table showing the number of cells before or after filtering based on the criteria written in Methods. **f**, UMAP embedding of snATAC-seq cells from *Enl*-WT kidneys. Cells are colored by annotated cell types. **g**, Genome browser view of the ATAC signal in each snATAC-seq cell type annotated in (**f**) at selected marker gene promoters. **h**, Heatmap showing the relative ATAC signal of 56258 cell type/lineage-specific peaks among all cell types annotated in (**f**). See Supplementary Table 5. ATAC signals were normalized by column Z-score. **i**, Table showing the top 2 most significant TF candidates from the motif analysis for cell type-specific peaks identified in (**h**). **j**, Heatmap showing the expression level of the TF candidates listed in (**h**) in the corresponding *Enl*-WT scRNA-seq dataset. The gene expression was normalized by row Z-score.

**Figure S6.**
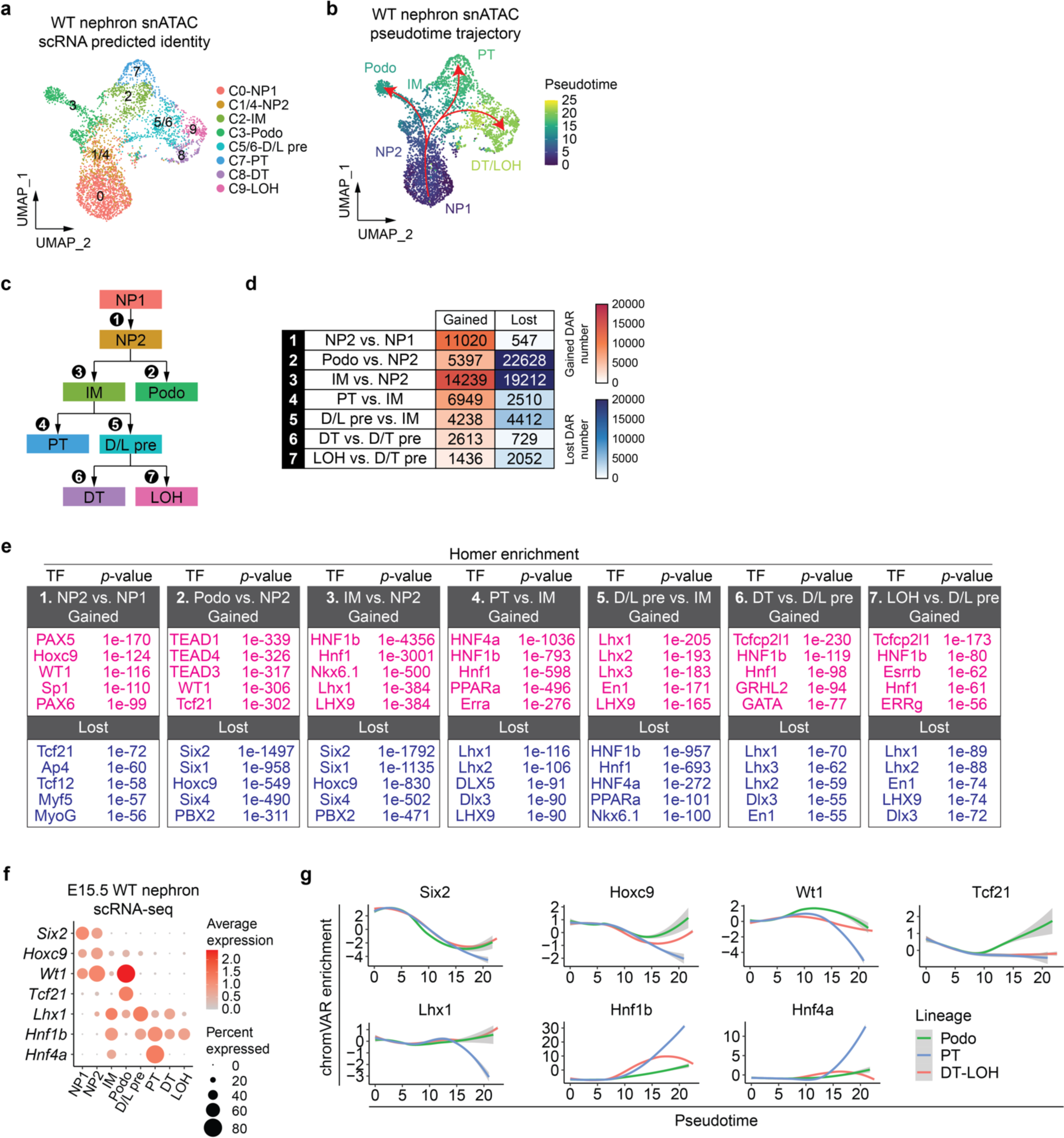
snATAC-seq reveals chromatin accessibility dynamics and the regulatory landscape during early nephrogenesis. **a**, UMAP embedding of snATAC-seq cells from *Enl*-WT nephrons. Cells are colored and labeled by the cell types predicted by corresponding scRNA-seq data. **b**, UMAP embedding of *Enl*-WT snATAC-seq nephron differentiation trajectory. Cells are colored by pseudotime. Trajectories are depicted by red arrows. **c**, Di-graph representing cell type divergence of *Enl*-WT nephrons. **d**, Table showing the number of differentially accessible regions (DARs) between subsequent stages in nephrogenesis. Cells were colored by the DAR number. Gained DARs were colored in red, lost DARs were colored in blue. See Supplementary Table 6. **e**, Table listing the top 5 most significant TF candidates from the motif analyses for the DARs identified in (**d**). TF candidates for gained and lost DARs were colored in red or blue, respectively. **f**, Dot plot showing the expression of selected TF candidates from (**e**) for each nephron cell type. Color scale represents the average expression level. Circle size represents the percentage of cells expressing the gene. **g**, Pseudotime-dependent dynamics of chromVAR TF enrichment score along Podo (green), PT (blue), and DT/LOH (red) differentiating trajectories.

**Figure S7.**
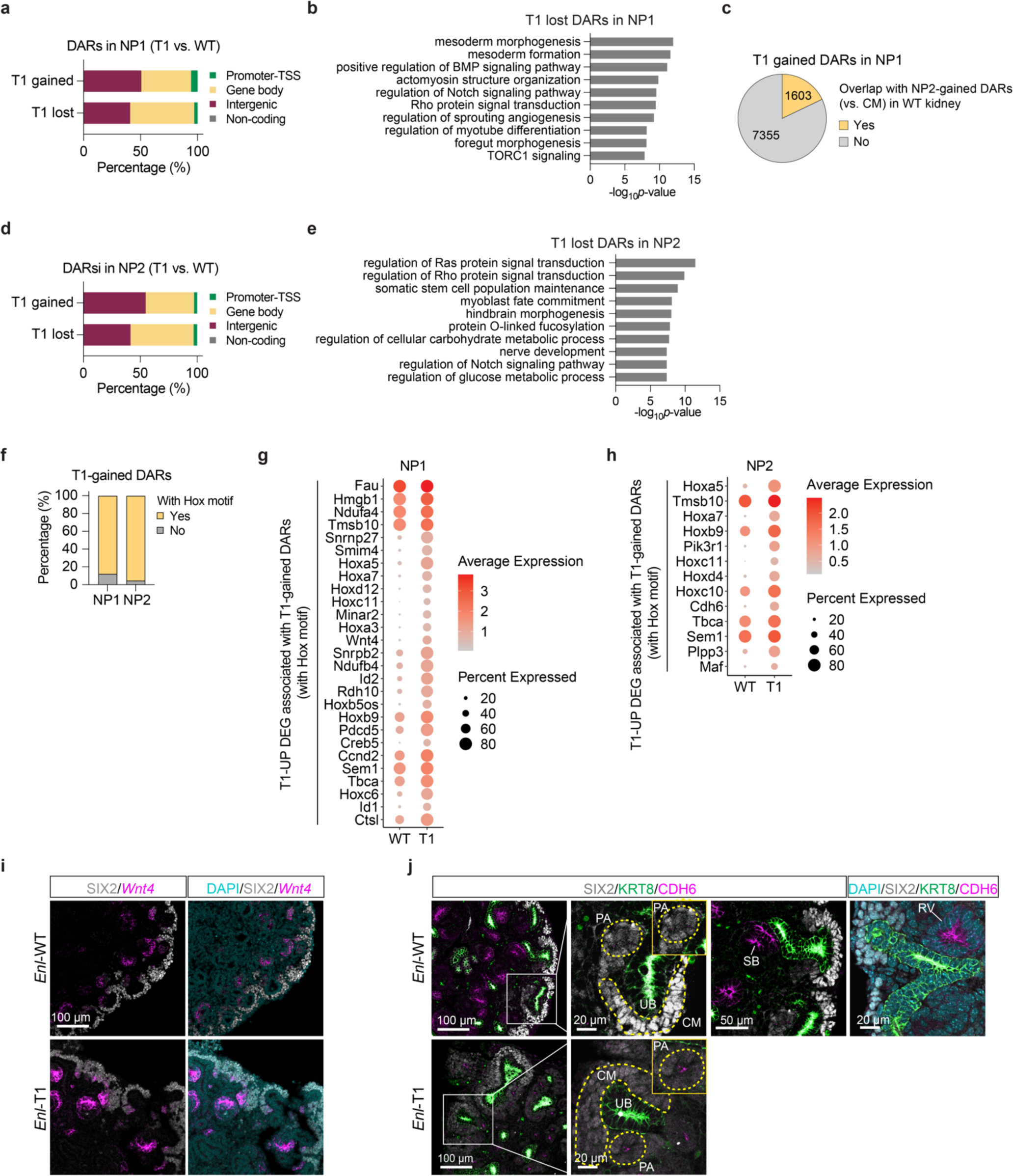
Mutant ENL promotes premature commitment of nephron progenitors while simultaneously restricting their differentiation through misregulation of specific TF regulons. **a**, **d**, Stacked bar plot showing the percentage of DARs located in promoter-TSS, gene body, intergenic, or non-coding regions. DARs were identified in NP1 (**a**) or NP2 (**d**) between *Enl*-WT and T1 cells. **b**, **e**, GREAT analysis of T1-lost DARs in NP1 (**b**) and NP2 (**e**). **c**, Pie chart showing the number of T1-gained DARs in NP1 overlapping with NP2-gained DARs (versus NP1) identified in *Enl*-WT kidneys. See Supplementary Table 8. **f**, Stacked bar plot showing the percentage of T1-gained DARs enriched with Hox TF motifs in NP1 (left) and NP2 (right). **g**, **h**, Dot plots showing the gene expression of Hox motif enriched T1-gained DARs associated T1-UP DEGs in NP1 (**g**) or NP2 (**h**). Color scale represents the average expression level. Circle size represents the percentage of cells expressing the gene. **i**, Representative images of SIX2/*Wnt4* mRNA co-staining in E15.5 kidneys. **j**, Representative images of SIX2/KRT8/CDH6 co-staining in E15.5 kidneys. CM, cap mesenchyme; UB, ureteric bud; PA, peritubular aggregate. SB, S-shape body.

**Figure S8.**
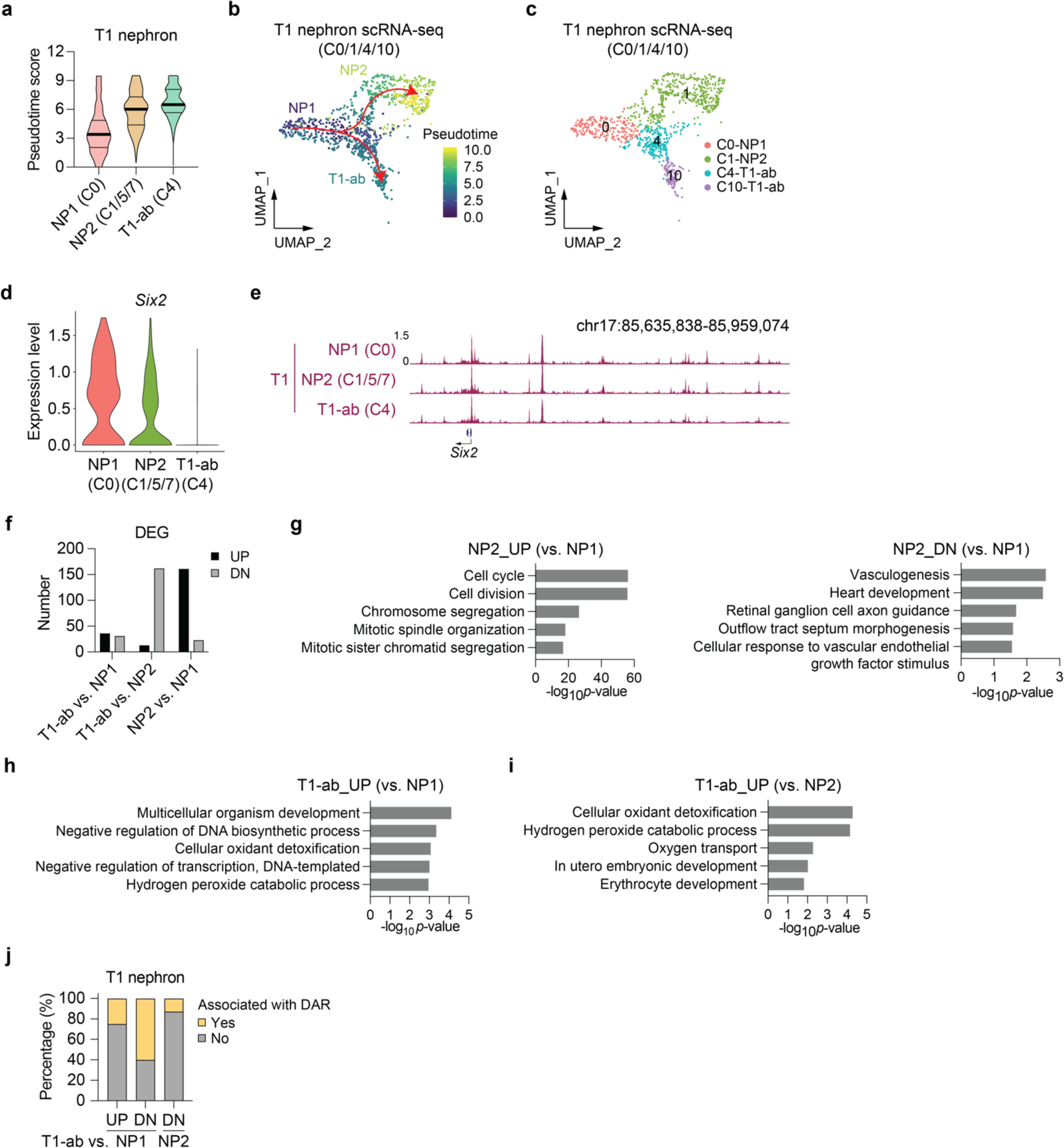
An abnormal progenitor state losing nephron chromatin identity emerges in the *Enl*-mutant kidney. **a**, Violin plot showing pseudotime score of *Enl*-T1 cells clustered in NP1, NP2, and T1-ab. Thick center lines indicate median, top and bottom line limits are set to the 25th and 75th percentiles. **b**, UMAP embedding of scRNA-seq *ENL*-T1 nephron differentiation trajectory from NP1 to NP2 or T1-ab, colored by pseudotime score. Trajectories are depicted by red arrows. **c**, UMAP embeddings of *Enl*-T1 nephron (NP1, NP2, and T1-ab) scRNA-seq differentiation trajectory (**b**) and cell clustering (**c**). Cells were colored by pseudotime score (**b**) or cell type (**c**). **d**, Violin plot showing the expression level of *Six2* in indicated cell types from *Enl*-T1 nephrons. **e**, Genome browser view of ATAC signal at *Six2* locus in indicated cell types from *Enl*-T1 nephrons. **f**, Bar plot showing the number of DEGs identified in indicated comparison. Upregulated (UP) and down-regulated (DN) DEG numbers were colored in black or grey, respectively. See Supplementary Tables 10. **g-i**, Bar plot showing the GO term analysis for the indicated DEGs. **j**, Stacked bar plot showing the percentage of DEGs associated with DARs identified in Figure 5b and c. See Supplementary Table 11.

**Figure S9.**
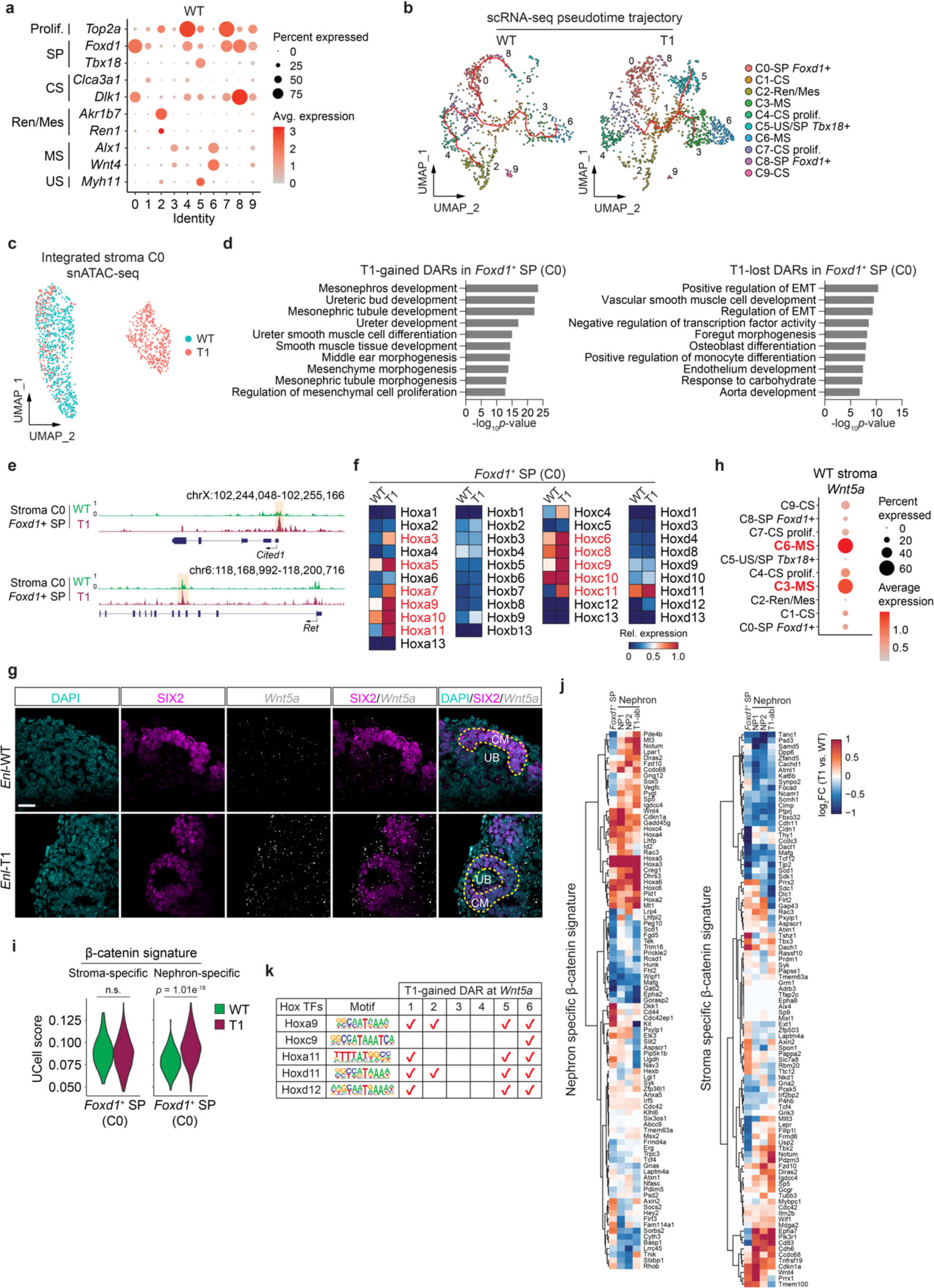
Figure 6. *Enl*-mutant *Foxd1*^+^ stromal progenitors exhibit altered chromatin accessibility and might affect stroma-nephron interactions through hyperactivation of the Wnt signaling pathway. **a**, The gene expression of cell type-specific marker genes in indicated cell clusters from *Enl*-WT stroma. Color scale represents the average expression level. Circle size represents the percentage of cells expressing the gene. SP, stromal progenitor; Prolif., proliferating cells; CS, cortical stroma; MS, medullary stroma; US, ureteric stroma; Ren/Mes, renin/mesangial cells; EMT, epithelial-mesenchymal transformation. **b**, UMAP embedding of *Enl*-WT (left) and T1 (right) scRNA-seq stroma differentiation trajectory. Cells were colored and labeled by cell types. Trajectories are depicted by red arrows. **c**, UMAP embedding of integrated snATAC-seq cells from *Enl*-WT and T1 stroma C0. Cells were colored by samples. **d**, GO term analyses for T1-gained (left) or lost (right) DARs in *Foxd1*^+^ SP cells identified in Figure 6f. **e**, ATAC signal at *Cited1* (top) and *Ret* (bottom) gene loci in *Enl*-WT and T1 stroma C0 cells. **f**, The expression levels of all 39 *Hox* genes in *Enl*-WT and T1 stroma C0 cells. **g**, Representative images of SIX2/*Wnt5a* mRNA co-staining in E15.5 kidneys. Scale bar = 20 µm. **h**, *Wnt5a* gene expression in each cell type of *Enl*-WT stroma. Color scale represents the average expression level. Circle size represents the percentage of cells expressing the gene. Two MS clusters with the highest *Wnt5a* expression level were highlighted in red. **i**, The UCell score evaluated by stroma (left) or nephron (right) specific β-catenin signature for cells from *Enl*-WT and T1 stroma. Wilcoxon rank-sum test *p*-values are shown. **j**, The fold change of expression level for the genes in nephron (left) or stroma (right) specific β-catenin signatures in indicated cell types. Fold change was scaled by log2. **k**, The enrichment of indicated Hox TF motifs within the T1-gained DARs in the *Wnt5a* gene locus.

**Figure S10.**
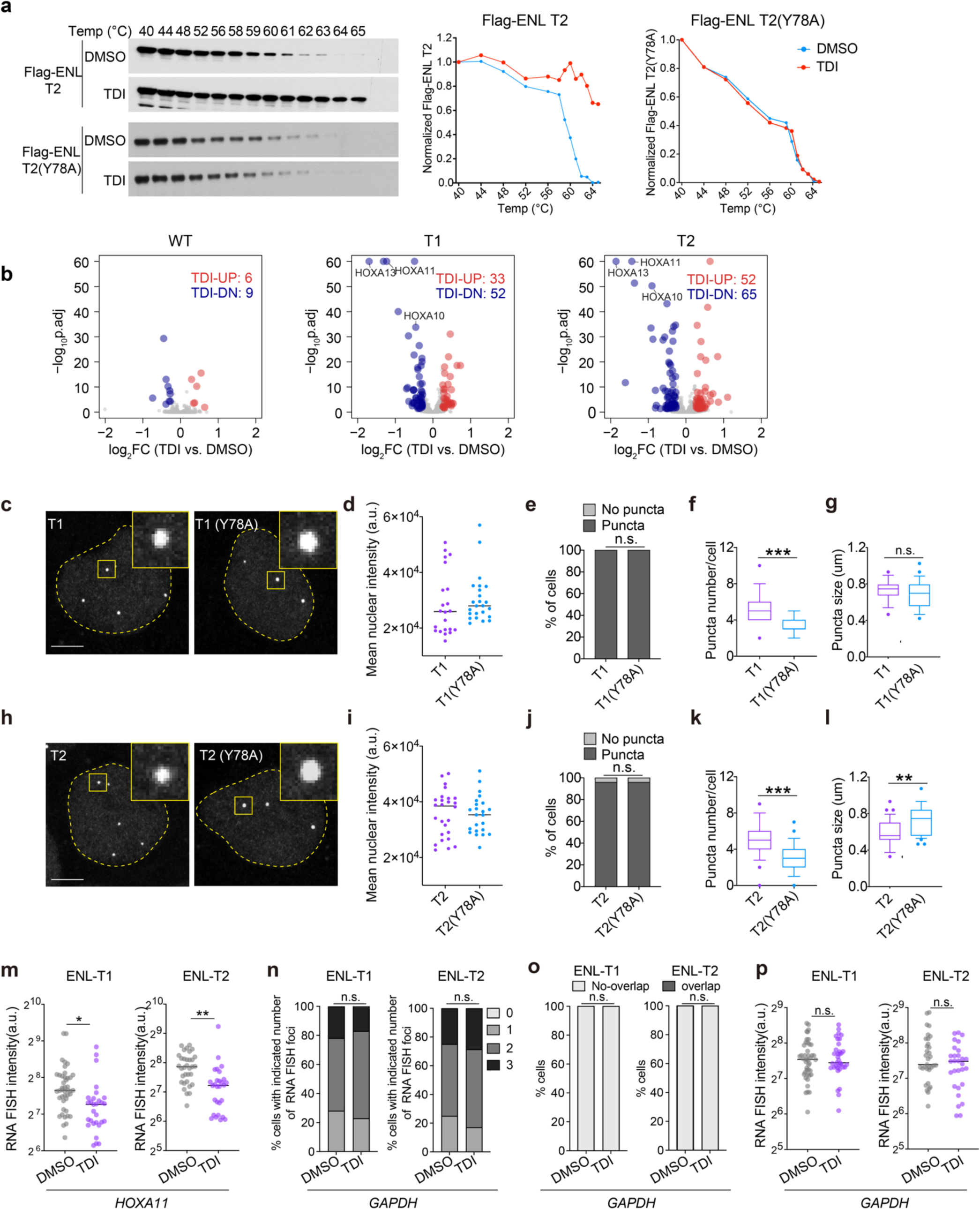
Blocking the acyl-binding activity of mutant ENL compromises its function on chromatin. **a**, Immunoblots and quantification showing the levels of Flag-ENL T2 and Flag-ENL T2 (Y78A) after heat treatment in HEK293 cells at increasing temperatures. **b**, Volcano plot of RNA-seq data obtained from HEK293 cells expressing near endogenous levels of indicated transgenes with DMSO or TDI treatment. Data represent the mean across three replicates. See Supplementary Table 15. **c**-**k**, IF staining of Flag-ENL T1 and T1(Y78A) (**c**) and Flag-ENL T2 and T2(Y78A) (**h**) in HEK293 cells. Scale bar, 10 µm. Quantification of the mean nuclear intensity (**d**, **i**), the percentage of cells with or without condensates (**e**, **j**), condensate number in each nucleus (**f**, **k**) and condensate size (**g**, **l**). Center lines indicate median and box limits are set to the 25th and 75th percentiles. Quantifications performed on 27-44 cells. **m**, Quantification of the mean intensity of *HOXA11* FISH foci. Center lines indicate the median. **n**, Quantification showing the percentage of cells with the indicated number of *GAPDH* nascent RNA FISH foci (number indicates the number of RNA FISH foci in each nucleus). **o**, The percentage of cells containing *GAPDH* nascent RNA FISH foci overlapping with Flag-ENL condensates. Chi-square test; **m**, Quantification of the mean intensity of *GAPDH* FISH foci. Center lines indicate the median. **e**, **j**, Chi-square test. **f**, **g**, **k**, **i**, **m**, **p** Two-tailed unpaired Student’s *t*-test. ***p < 0.001, **p < 0.01, *p < 0.05, n.s., no significance

**Figure S11.**
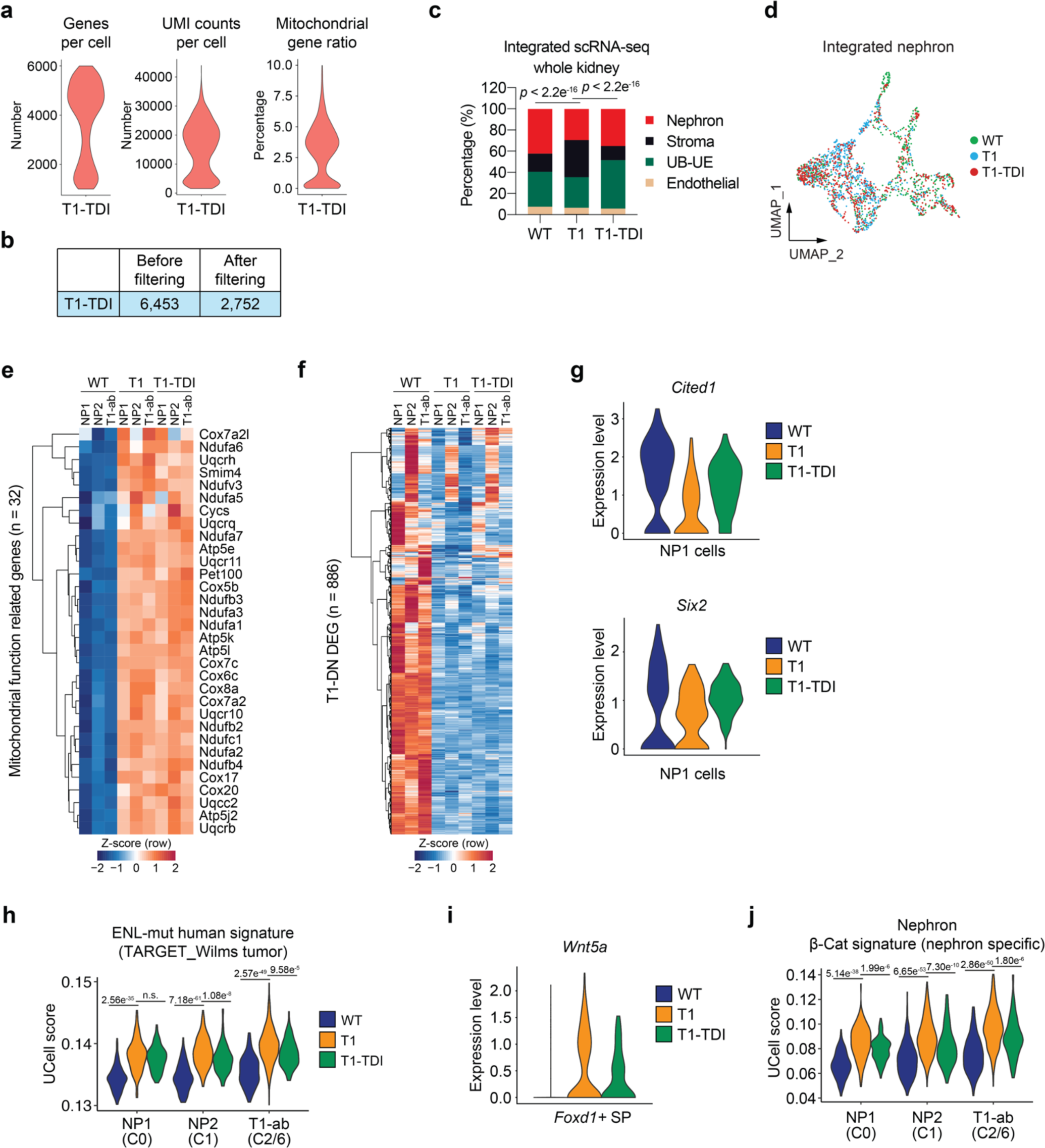
Transient *in vivo* treatment with TDI-11055 partially rescues mutant ENL-induced developmental and transcriptional defects in the developing kidney. **a**, Violin plot showing the number of informative genes per single cell, unique molecular identifiers (UMIs) per single cell, and mitochondrial gene ratio in scRNA-seq dataset of *Enl*-T1 kidneys with TDI-11055 treatment (T1-TDI) after quality control filtering. **b**, Table showing the number of cells before and after filtering according to the criteria written in Methods. **c**, The percentage of four main embryonic kidney compartments within samples. Chi-Square test *p*-value is shown. **d**, UMAP embedding of integrated scRNA-seq data from *Enl*-WT/T1/T1-TDI nephrons. Cells were colored by sample. **e, f**, Heatmap showing the fold change of mitochondrial and metabolism-related T1-UP DEGs identified in Figure 2i (**e**) and union T1-DN DEGs identified in Figure 2m **(f)** within the indicated nephron cell types. Fold change is scaled by row z-score. **g**, Violin plot showing expression level of *Cited1* (top) and *Six2* (bottom) in NP1 from *Enl*-WT, T1, and T1-TDI nephrons. **h**, Violin plot showing the UCell score evaluated by human ENL-mut signature for the NP1, NP2, and T1-ab cell types from *Enl*-WT, T1, and T1-TDI nephrons. Wilcoxon rank-sum test *p*-values are shown. **i**, Violin plot showing expression level of *Wnt5a* in *Foxd1*^+^ SP from *Enl*-WT, T1, and T1-TDI stroma. **j**, Violin plot showing the UCell score evaluated by nephron specific β-catenin signature for the NP1, NP2, and T1-ab cell types from *Enl*-WT, T1, and T1-TDI nephrons. Wilcoxon rank-sum test *p*-values are shown.

